# Multi-Domain Interplay Controls Full-Length TDP-43 Phase Separation and Condensate Dynamics

**DOI:** 10.64898/2026.06.03.729912

**Authors:** Xiaofei Ping, Rodrique G. M. Badr, Saskia Hutten, Xinxiang Chen, Lucia Baltz, Mahesh Yadav, Jana Sophie Höppner, Friederike Schmid, Dorothee Dormann, Lukas S. Stelzl

## Abstract

TDP-43 (TAR DNA-binding protein 43) is a 414-amino acid protein with a structured N-terminal domain (NTD), two RNA recognition motifs (RRM1 and RRM2), and a long disordered C-terminal low complexity domain (LCD). TDP-43 forms phase-separated condensates as part of its physiological function in RNA processing. However, aberrant TDP-43 condensation, changes in condensate material properties, and subsequent aggregation are linked to the development of neurodegenerative diseases. Using explicit-solvent, near-atomic resolution coarse-grained simulations we demonstrate how the interplay among different domains drives phase separation of the full-length protein.We directly capture how the secondary structure of a conserved helix in the LCD modulates phase separation, and follow the effect of phosphomimicking mutations on the condensation of full-length TDP-43. C-terminal phosphomimicking mutations rewire the interactions of the LCD by increasing solvation locally and enhancing Na^+^ binding to the LCD. Our simulations and *in vitro* experiments emphasize the importance of the aromatic residues in the LCD but also of N-terminal residues 1-101 including the NTD for full-length TDP-43 condensation. With a Gō-type approach we capture not just conformational flexibility but also specific dimer formation through NTD-NTD interactions and how they modulate the phase behavior as well as the dynamic and interfacial properties of full-length TDP-43 condensates.

## INTRODUCTION

TAR DNA binding protein of 43 kDa (TDP-43) is a pathological hallmark of aggregates found in the brains of patients suffering from frontotemporal dementia (FTD), amyotrophic lateral sclerosis (ALS), limbic-predominant age-related TDP-43 encephalopathy (LATE) as well as Alzheimer’s disease^1–3^. As an RNA-binding protein (RBP) with a central role in RNA processing, TDP-43 localizes mainly in the nucleus^4,5^ but in disease is progressively lost from the nucleus and instead accumulates in cytosolic inclusions of neurons and glial cells^6,7^. TDP-43 can undergo phase separation in cells^8,9^, where liquid TDP-43 condensates are important for the proper functioning of neurons, whereas more viscous condensates or solid-like assemblies impair neuronal physiology^10^.

TDP-43 is a 414–amino acid protein comprising three folded domains, namely the N-terminal domain (NTD, aa 1–80) and two RNA recognition motifs domains (RRM1, aa 105–176 and RRM2, aa 192–259), along with a long C-terminal intrinsically disordered region (IDR) and a nuclear localization signal (NLS) sequence that links the NTD to RRM1. Within the C-terminal IDR, also known as the low-complexity domain (LCD), a conserved region (CR) (aa 319–341) contains a short segment (aa 320–332) with a high propensity to form an *α*-helix. This helical element has been identified as a critical structural determinant of liquid–liquid phase separation^9,11–15^, and most structural and biophysical studies of this helical region have focused on LCD constructs.

The NTD of a TDP-43 molecule can bind with the NTD of another TDP-43 molecule to form dimers. NTD-dimerization is essential for the phase separation of full-length TDP-43 and contributes to its biological function^16–19^. Loss of TDP-43 dimerization is associated with neurodegenerative disease^20^. Meanwhile, C-terminal fragments comprising the LCD plus portion of the RRM are often found in patients^21–24^ hence the LCD has been often studied as a proxy to understand disease-linked TDP-43 phase separation^12,25,26^. While the LCD itself can undergo phase separation by itself^11,25,26^, the difference in phase behavior of full-length TDP-43 and the isolated LCD has not been characterized in detail.

Post-translational modifications such as phosphorylation, have emerged as important regulators of phase separation, especially for RBPs^27,28^. Aggregated TDP-43 isolated from patient samples is hyperphosphorylated in its C-terminal LCD, and the presence of specific phosphorylation sites is commonly used as a pathogenic marker^29–33^. However, recent studies suggest that C-terminal hyperphosporylation of TDP-43 may actually serve as a cytoprotective mechanism as it increases TDP-43 solubility both *in vitro* and in cells and decreased its tendency to undergo phase separation and aggregation^8^. In multi-scale molecular dynamics simulations of LCD condensate, we could recently show that phosphomimetic mutations result in increased solvation and hence reduced homotypic interactions^8^. Whether similar effects arising from protein–solvent interactions and electrostatic repulsion^34^ between charged residues also apply to full-length TDP-43 remains an open question that requires further simulations.

Advances in coarse-grained (CG) modeling enable simulations of multi-domain proteins containing both structured and intrinsically disordered regions with near-atomic detail^35–38^. Mohanty *et al*. previously assessed the interactions of different regions of TDP-43 in residue-level coarse-grained simulations^39^. More-over, near-atomic-resolution coarse-grained simulations with explicit solvent^40–42^ can capture the phase separation of full-length proteins including TDP-43^43^. The Martini3 coarse-grained force field enables simulations that reveal specific interactions of individual residues with biomolecules, ions, and solvent molecules. Thomasen *et al*. demonstrated that such near-atomic-resolution coarse-grained simulations can capture the effects of mutations on the global conformations of IDR-rich proteins^44,45^. Other studies have shown that Martini3 simulations can also reproduce the effects of IDR truncations on phase separation^41^. However, it remains unclear whether coarse-grained simulations can accurately recapitulate the effects of smaller sequence changes on phase behavior. Another challenge in coarse-grained simulations is capturing the conformational dynamics of folded domains. Simulations have been employed to understand how inter-chain interactions of disordered regions and the interplay of disordered regions^46,47^ determine the material and dynamic properties of phase-separated condensates^48^. To preserve the initial folded structure, the Elastic Network (EN) approach can be employed in CG simulations by applying harmonic restraints on native contacts, thereby preventing their dissociation within folded regions^49^. Meanwhile, conformational malleability can be captured by Gō-type approaches^50–52^ instead of EN. Native contacts in Gō-type approaches are represented by Lennard-Jones (LJ) potentials and can dissociate when the interaction is not sufficiently stabilizing. Furthermore, dimerization and multimerization can in principle be modeled by extending Gō-type approaches to intermolecular interactions, as has been demonstrated for amyloid fibril formation^53^; however, extensions to inter-chain interactions of folded protein domains in protein condensates are still lacking.

Here, we use coarse-grained molecular dynamics simulations with near-atomic detail to provide molecular-level insights into phase-separated full-length TDP-43. Multiple simulations with durations beyond 20 *µ*s (including wild-type (WT) and variants, each with replicas; approximately 400 *µ*s in total) reveal the effects of point mutations and truncations and are consistent with experiments using recombinant proteins *in vitro*. The results provide molecular-scale insight into how C-terminal phosphomimetic mutations and mutations of key aromatic residues (phenylalanines, tryptophans) in the LCD reduce phase separation propensity of full-length TDP-43. Furthermore, our simulations reproduce the enhanced phase separation propensity of full-length TDP-43 compared to the isolated LCD. Additionally, we show how to capture the NTD-mediated dimerization of TDP-43 and suggest that inter-chain interactions of the NTD and the adjacent NLS region contribute to the condensation behavior. In summary, our simulations show that phase separation of full-length TDP-43 is governed by a combination of homotypic and heterotypic inter-chain interactions involving multiple regions of the protein, in addition to the well-established contributions of the helical region and the aromatic and aliphatic residues in the LCD.

## RESULTS

### Phase behavior of full-length TDP-43 in chemically-detailed coarse-grained molecular dynamics simulations

To characterize the phase behavior of full-length TDP-43 and the molecular drivers of its phase separation, we performed simulations with the transferable and near-atomic resolution coarse-grained Martini3 model^40^. To investigate the effects of varying solution conditions, we varied the protein-water interaction strength through a rescaling factor *λ*^44,45,47,54^. Reducing the value of the scaling factor *λ* mimics the addition of crowding agents or a decrease in temperature, while increasing *λ* corresponds to a removal of crowding agents or an increase in temperature. The original Martini3 model interaction corresponds to *λ* = 1. In previous work on the simulation of multi-domain proteins, setting *λ* = 1.06 yielded a good agreement between simulations and experiments^45^. Following this, unless otherwise specified, we use *λ* = 1.06 as our default value. Fig 1A shows an atomic resolution structure of a single full-length TDP-43 chain, while Fig 1B shows full-length TDP-43 as represented by the Martini3 EN model^49^ with enhanced protein-water interactions using *λ* = 1.06^45^. Coarse-grained simulations of single chain TDP-43 capture its global dimensions when compared to solution scattering experiments^55,56^ (Fig. S1 and Fig. S2) in addition to its conformational flexibility as determined by atomistic molecular dynamics simulations (Fig S3). Movie S1 illustrates how a solution at a protein concentration of 2.77 mM (70 chains) phase separates to form a protein droplet in a cube. Fig 1C, D show the starting and final configuration of the system, respectively. To quantify the phase behavior of TDP-43 we performed simulations with 60 chains in a slab geometry (Fig. S4, Fig. S5), which leads to faster equilibration of the system^57^. Tracking the protein concentration profiles along z-axis over time^58^ for *λ* = 1.06 (Fig 1F) and Fig. S5, Fig. S6) reveals that TDP-43 chains in the condensates can move to the dilute phase on the simulation time scale of tens of *µs*. Lowering *λ* to 1.05 from our default value of 1.06, thus weakening the protein-water interactions, reduces the number of proteins in the dilute phase, as can be seen from the time-dependent concentration shown in Fig 1E. Chains tend to stay in the condensate (Fig 1E and H). Meanwhile, strengthening the protein-water interactions using *λ* = 1.07 destabilizes the condensate, as evidenced by a lower concentrations of the dense phase and many chains in the dilute phase (Fig 1G and J and Fig. S6).

**FIG. 1.**
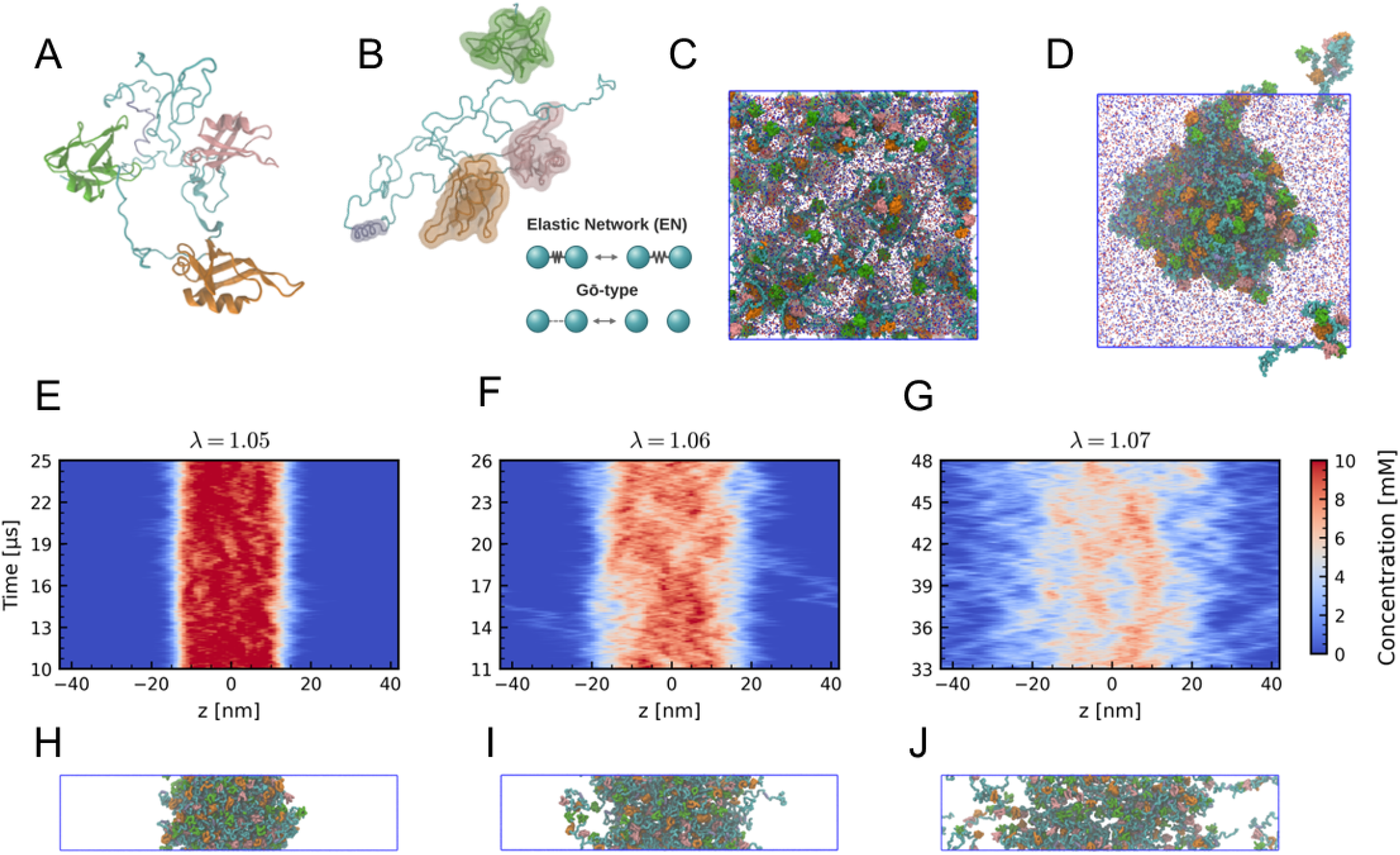
Phase separation of full-length TDP-43 in coarse-grained molecular dynamics simulations. (A-B). Representative structures of (A) all-atom (AA) and (B) Martini3 coarse-grained (CG) molecular dynamics simulation of full-length protein TDP43. The represented CG model is implemented with Elastic-Network (EN) for the folded regions, but Gō-type approaches can capture dynamics beyond breathing motions. Residues highlighted in green, orange, pink correspond to the N-terminal domain (NTD), RNA-recognition motif 1 (RRM1), and 2 (RRM2), respectively. Cyan areas indicate the protein’s disordered regions (IDR), and the region colored in purple represents the helical segment within the C-terminal IDR. Water and ions are omitted for clarity. (C) Initial configuration of the Martini3 EN model for TDP-43 with protein-water interaction strength *λ* = 1.06 in a cubic simulation box. (D) Representative snapshot of running the simulation with *λ* = 1.06 for the starting configuration shown in C. The proteins form a liquid-like spherical droplet in the cubic box. Also shown are Na^+^ (red) and Cl^−^ (blue) ions. Water beads are omitted for clarity. (E-G) Time-dependent concentration profiles for simulations for simulation with the Martini3 EN model with different protein-water interaction strength *λ* = 1.05, *λ* = 1.06, and *λ* = 1.07. The protein concentration is indicated by the color bar. (H-J) Snapshots present the protein condensate in the Martini3 EN model with protein-water interaction strength *λ* = 1.05, *λ* = 1.06, and *λ* = 1.07 respectively along the z-axis.

To identify contacts/interactions that drive full-length TDP-43 phase separation in our simulations, we searched for prominent inter-chain contacts. Notably, the LCD forms extensive inter-chain contacts, particularly with the LCD regions of neighboring chains (Fig 2A). Fig 2B shows an example of how the LCD regions of two TDP-43 molecules can interact within a condensate. The helix within the CR frequently engages in interactions with the same region (aa 320-332) on adjacent chains (Fig 2A). Aromatic residues within the C-terminus play a prominent role in mediating these inter-chain contacts (Fig 2A and D). In addition, the LCD engages in numerous interactions with the RRM2 domain (Fig 2C), many of which involve the conserved helical region (Fig 2A). Furthermore, the LCD also interacts with the NLS region which connects the folded NTD to the RRM1 (Fig 2A). The C-terminal part of the LCD features many interaction-prone residues, in particular aromatic residues; overall, however, contacts are distributed along the entire protein sequence (Fig 2D). For example, two highly interacting aromatic residues (W113 and W172) are within the folded RRM1 domain. The three folded domains NTD, RRM1, RRM2 show relatively modest self-interactions, but interact mainly with the LCD (Fig 2A). In summary, all domains feature residues that engage in few inter-chains interactions, alongside highly interaction-prone residues such as aromatic residues. In our analysis the LCD stands as its aromatic residues and the CR helix form many contacts with other chains in our simulations (Fig. 2A).

**FIG. 2.**
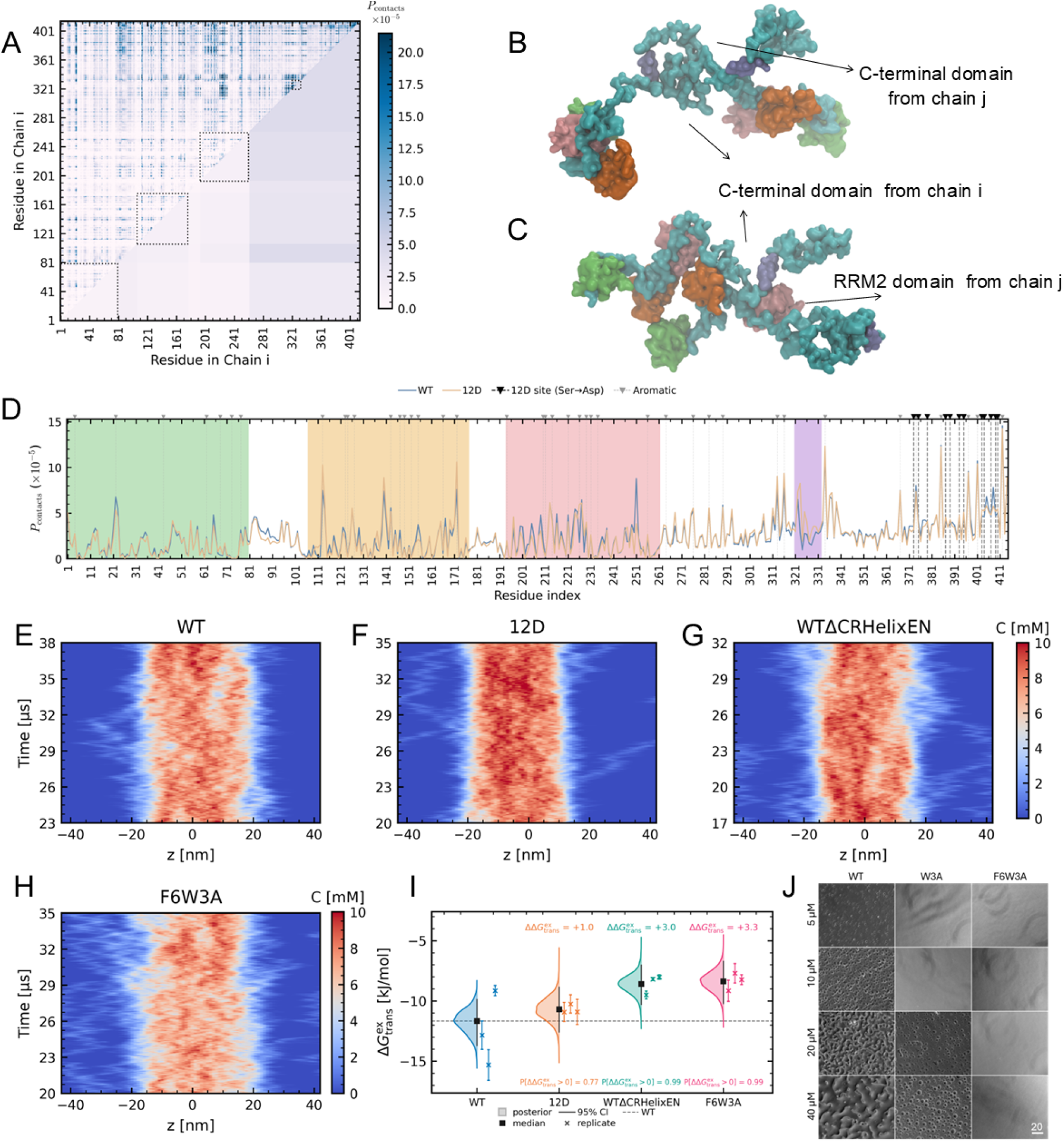
Sequence-specific interactions of full-length TDP-43 in condensates. (A) Inter-chain contacts mapped at *λ* = 1.06 in EN model. Dashed rectangles indicate structured regions with restraints. The upper triangle shows residue-wise contact map, the lower triangle displays domain-wise contact map by summing up the contact possibility and averaged by the total residue number within the specific domain, color bar on the right represents the residue contact probabilities. (B-C) Snapshots illustrating LCD–LCD (B) and LCD-RRM2 (C) inter-chain contacts. (D). Averaged intermolecular contact probability for each residue shown for in WT (blue) and mutant 12D (yellow). Black arrows and dashed lines highlight the positions of serine residues that were replaced by aspartate residues, while gray arrows and dashed lines denote aromatic residues. (E-H) Time-dependent concentration histograms along the z-axis for the WT protein (E), mutant 12D (F), WTΔCRHelixEN (G), and mutant F6W3A (H), protein concentration indicated by the color bars on the right. (I) Excess transfer free energies of the mutants in slab-geometry simulations at *λ* = 1.06, 95% Bayesian Credible Interval of each group is indicated by shading, individual replicates are shown as colored circles; error bars denote standard errors propagated from dilute and dense phase concentration measurements. (J) *In vitro* phase-separation assays for TDP-43 WT and the aromatic mutants (W3A and F6W3A) show diminished condensate formation compared to the WT for all tested concentrations. Scale bar: 20 *µ*m.

### Chemically-detailed coarse-grained simulations reveal how mutations affect TDP-43 phase behavior

Next, we sought to test if we can resolve the impact of sequences changes on the phase behavior of full-length TDP-43 by coarse-grained simulations. In particular, we examined how the phosphomimetic 12D variant, in which twelve serine residues in the TDP-43 C-terminus were substituted with aspartic acid to mimic disease-linked C-terminal phosphorylation^8^, reduces TDP-43’s propensity to phase separate. Each variant was simulated using three independent replicates, with the same conditions (about 300,000 beads in total including about 54000 protein beads) and simulation parameters as in simulations of WT. Simulations over 30 *µ*s enable the quantification of how the 12D mutant affects protein concentrations in both the dense and dilute phases. Similarly to WT TDP-43, an equilibrium between two phases is established for the 12D variant. Proteins continuously exchange between condensates and the surrounding solution, as shown by the time-dependent concentration profiles in Fig 2F, Fig. S8, and the dilute phase concentration in Fig. S9. Consistent with experimental observations^8^, the 12D mutant exhibits a reduced propensity for phase separation, which we quantified with a Bayesian model (see Methods for details). In the simulation with 12D TDP-43 more chains are found in the dilute phase than for WT TDP-43 (Fig 2F, Fig S8 and Fig. S9), but the concentration of the dense phase is also higher than for WT (Fig. 2F, Fig. S7 and Fig. S10). The excess transfer free energy 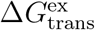 (Eq. 3) and Supporting Information) is less negative for the 12D mutant than for WT indicating reduced phase separation with a probability 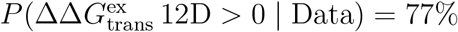. This reduction is associated with a loss of intermolecular contacts in the distal C-terminal region of the LCD (Fig 2D), particularly near residues S403, S404, S407, S409, and S410. The increased net negative charge of −16 for the 12D variant instead of −4 for WT leads to enhanced recruitment of Na^+^ ions to the condensate (Fig S11). 12D TDP-43 chains are surrounded by Na^+^ both in the condensate and in the dilute solution and the binding of counter ions might make it more favorable for 12D chains to be outside the condensate in the dilute phase and, contributing to an increase in *c*_dilute_ compared to WT TDP-43. Besides compensating for the negative charge of the Asp groups, Na^+^ ions may also bridge between Asp groups. Such ion-mediated contacts and new protein contacts of Asp at the phospho sites may explain the increased concentration of the dense phase for the 12D variant (Fig. S12). At the sites of the phosphomimetic mutations, inter-chain contacts are re-wired compared to WT, as the surrounding region is no longer neutral as it was in the WT protein. Consequently, the number of contacts of the phospho sites with the C-terminus of the LCD drops. While interactions with, e.g., negatively charged partner residues are lost (Fig. S12)A), new interactions with Lys and Arg are formed, but the Asp residues at the phospho-sites also engage in new interactions with Asn and Gln (Fig. S12)A). At the sites of the phosphomimetic mutations, more water is bound to Asp rather than Glu residues, which is in line with previous all-atom simulations of the WT and 12D LCD condensates^8^ (Fig. S12B).

To support the role of aromatic residues identified in our simulations (Fig 2A), we performed phase separation assays *in vitro* using recombinant purified TDP-43. For this, we compared the phase separation propensity of TDP-43 and TDP-43 mutants in which specific aromatic residues had been mutated to alanine, as previously described^26,59^. Consistent with the findings for the isolated TDP-43 LCD^26,59^, we observe that full-length TDP-43 W3A (W334A, W385A and W412A) carrying three W-to-A substitutions, as well as a F6W3A mutant carrying six additional substitutions of F-to-A (F276/283/289/367/397/401^60^) show strongly impaired condensation propensities compared to TDP-43 WT. W3A forms few, small condensates at very high protein concentration of 40 *µM*, while for F6W3A no condensates are visible even at this concentration (Fig 2J). Previously work has already demonstrated that certain point mutations in the CR region, like the exchange of the W in the 334th position into A, reduced the helicity of this region^12^ At the same time, the W334A mutation reduced the ability of TDP-43 to phase separate^12^. Since our W3A and F6W3A mutants above both include the W334A mutation, we investigated the role played by the secondary structure versus the role of aromatic residues in facilitating phase separation.

To disentangle the effect of losing inter-chain contacts mediated directly by W and F residues and the effects inferred by the destabilization of the C-terminal helix in the CR region^12^, we performed simulations where we removed the EN restraints that support the helical structure of amino acids 320-332 (WTΔCRHelixEN). Removing the EN restraints in WT TDP-43 reduces its ability to phase separate with a probability 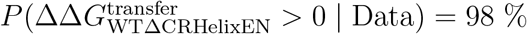 (Fig 2I). As a result of the loss of structure in the CR, we observe more chains outside the condensate (Fig 2G, I, Fig. S7, Fig S8 and Fig. S9) while the dense phase concentration actually increases, likely as the packing restraints due the folded structure of the CR helix are lost (Fig S10). Exchanging residue types according to the F6W3A mutant in addition to having a free helix leads to a slightly less dense condensate compared to both WT and WTΔCRHelixEN, as evident by a reduction in *c*_dense_ (Fig S10B). However, within our simulation time, we do not observe a significant difference in the number of chains in the dilute phase between the and WTΔCRHelixEN and the F6W3A free helix variants (Fig S10A). Simulation durations beyond ≈ 30 *µs* are likely required to really establish any differences in the dilute phase concentration (Fig S8). Despite the slight reduction in the dense phase concentration, the F6W3A mutation with a free helix does not further reduce the propensity of TDP-43 to phase separate when compared to the WTΔCRHelixEN variant, with both showing very similar Δ*G*^transfer^ (Fig. 2I). Consistent with this, the F6W3A free helix variant has a probability 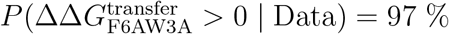 to have a reduced propensity compared to WT TDP-43. Importantly, at the sites of the F6W3A mutations, TDP-43 makes fewer contacts (Fig S13). Taken together, we show that we can use near-atomic resolution to directly track the impact of changes in secondary structure and changes to due to mutations of individual amino acids, which is challenging by experiment alone. Notably, we demonstrate that the secondary structure of the CR helix is a key driver of the phase separation of full-length TDP-43.

### Simulations and experiments quantify the role of the TDP-43 LCD in the phase separation of full-length TDP-43

Considering that multiple key intermolecular contacts in TDP-43 condensates are formed by the LCD, we sought to assess its importance for the phase separation of full-length (FL) TDP-43 by simulations and experiments. Considering that the LCD (154 aa, residues 261-414) is much shorter than full-length TDP-43 (414 aa), we wondered whether the LCD would phase separate under the same conditions as full-length TDP-43 phase separates. Indeed, LCD chains do not form condensates under the solvent conditions (*λ* = 1.06) and average number density of residues at which full-length TDP-43 readily phase separates (Fig 3A,B). At lower values of *λ* of 1.04 and 1.05, where solvent conditions more strongly favor condensation, LCD condensates are formed (Fig 3A,C). At *λ* = 1.05 the dense phase concentration of TDP-43 is sill considerably lower than full-length TDP-43, but at *λ* = 1.04, the dense phase concentration is 6 mM, close to the dense phase concentration of the full-length TDP-43. However, at *λ* = 1.06, preformed LCD condensates generated at *λ* = 1.04 dissolve over time, confirming that the solvent condition *λ* = 1.06 does not favor LCD condensates. Besides sequence length, full-length TDP-43 and LCD also differ in their charge profiles. A higher net charge per residue (NCPR) can reduce the propensity of a protein to phase separate^61^. However, the difference in NCPR between WT (NCPR −0.001) and LCD (+0.026) does not account for the difference, since the NCPR is lower than that of the 12D variant (−0.038), which phase separates much more strongly in our simulations. As a consequence of the larger NCPR, the LCD condensates binds more ions than WT TDP-43 condensates (Fig 3D). Experiments are in line with the results of our simulations, where we observe an apparently reduced phase separation of LCD compared to full-length TDP-43. In experiments, full-length TDP-43 droplets are abundant and clearly visible at a protein concentration of 5 *µ*M. At this protein concentration no droplets are detected for LCD. Increasing the protein concentration to 20 *µ*M, leads to the appearance of irregular amorphous structures under the microscope, which may suggest that the LCD starts to form condensates (Fig 3E). Together, this comparison shows that coarse-grained simulations with the Martini3 simulation model capture experimental trends in phase separation due to sequence truncations.

**FIG. 3.**
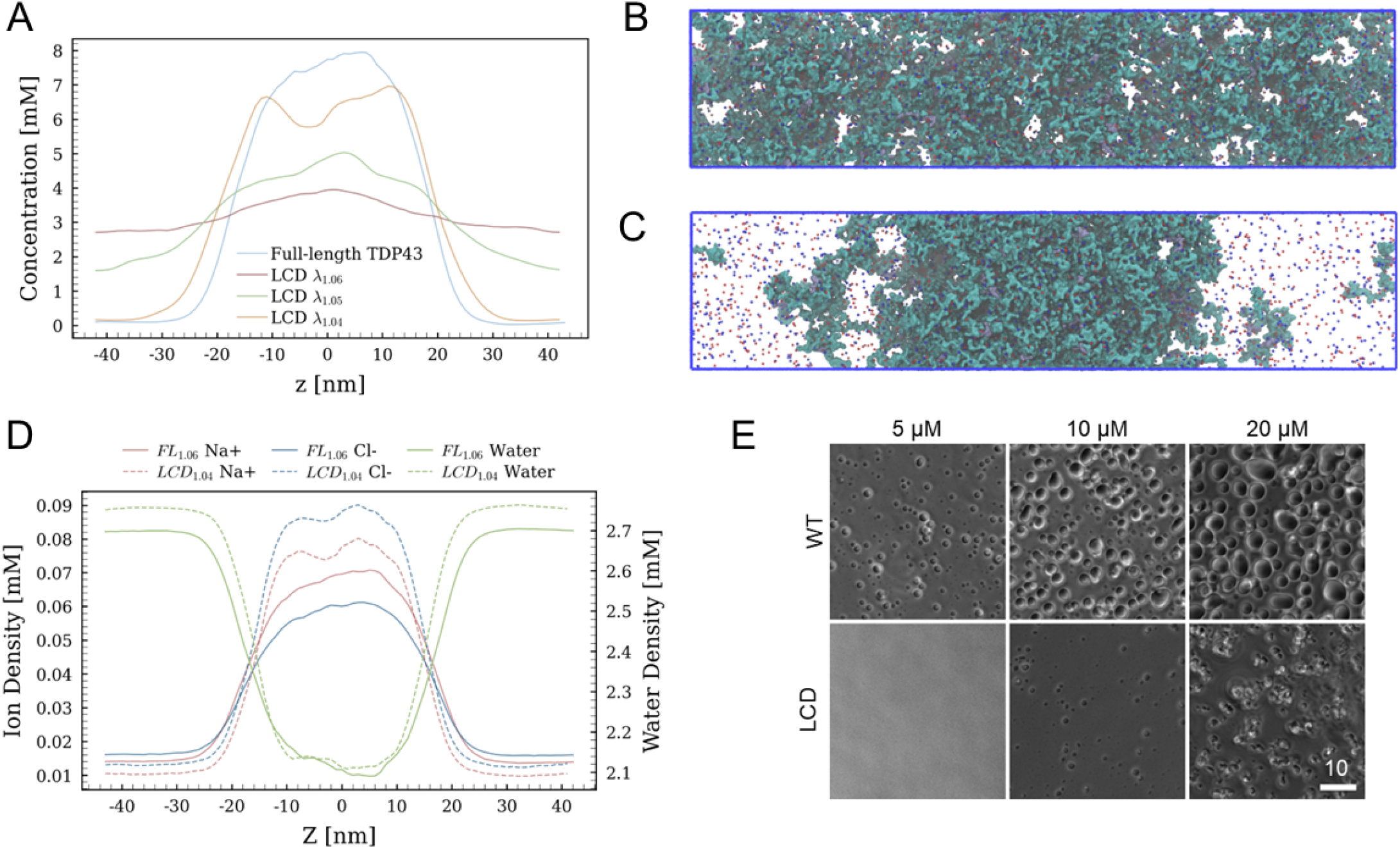
TDP-43 full-length (FL) and LCD phase separation in Martini 3 EN simulations. (A). Protein concentration profiles along the z-axis. Full-length TDP-43 was simulated with *λ* = 1.06, and LCD chains were simulated with *λ* = 1.06, *λ* = 1.05, and *λ* = 1.04 in the slab geometry. (B). Representative snapshot of full-length (FL) TDP-43 condensate at *λ* = 1.06. (C). Representative snapshot of LCD forming a condensate at *λ* = 1.04. (D). Ion and water density profiles along the z-axis for full-length TDP-43 (*λ* = 1.06) and LCD (*λ* = 1.04). (E) *In vitro* experiments testing phase separation of full-length TDP-43 and the isolated LCD at different protein concentrations.

To quantitatively understand the difference in phase behavior of full-length TDP-43 and LCD we determined their saturation concentration (*c*_sat_ = *c*_dilute_) as previously described^62^. For this, we quantified experimentally the protein concentration of TDP-43 FL or LCD in the dilute phase using a sedimentation assay followed by SYPRO Ruby staining (Fig. S14). The experimental *c*_dilute_ value for TDP-43 FL was substantially lower (0.51 ± 0.17 *µ*M) than that of the LCD alone (~ 4.5 ± 0.06 *µ*M). This is a further indication that the LCD-LCD interactions are not the sole drivers of phase separation. However, the reduction in phase separation propensity could be due to the difference in length between the two sequences. To argue against the possibility that this change is simply due to the difference in protein length, we refer to Flory-Huggins solution theory. From the Flory-Huggins model we can predict how the length of a homopolymer determines its *c*_dilute_ (Supporting Information). Fig. S15 shows that, for a homopolymer at fixed interaction strength, the expected change in *c*_dilute_ spans many orders of magnitude upon a change in length compared to TDP-43 FL versus LCD. Although Flory-Huggins theory applies to homopolymers, while proteins are (charged) heteropolymers, the relatively small difference in *c*_dilute_ cannot be explained on the grounds of length alone. This conclusion is in line with the many intermolecular interactions centered on the LCD (Fig. 2) that assist with phase separation, including the conserved helix, aliphatic and aromatic interactions^11,12,63^. Conversely, the net reduction in phase separation propensity highlights the importance of interactions of the LCD with other domains as well as the interaction of other domains among each other.

All in all, contact statistics in simulations and truncations in simulations and *in vitro* experiments, reveal the importance of the LCD for driving the phase separation of full-length TDP-43, but also suggest that other sequence elements beyond the LCD contribute to TDP-43’s phase behavior.

### Conformational flexibility of TDP-43 folded domains

To broaden our view of the molecular drivers of TDP-43 phase separation, we investigated how the folded domains contribute to TDP-43 condensation, considering both the conformational flexibility of the folded domains and the dimerization of NTD. First, we wondered how local structural flexibility, beyond the breathing motions captured by the EN, affect TDP-43 phase separation and found that increasing conformational flexibility can enhance phase separation (Fig. 4A). For this we employed a *Gō*_intra_ model, where the (Lennard-Jones) interactions that stabilize the folded domain structure can be broken unlike in an EN model. The *Gō*_intra_ bonds then allow the folded domains to deform and even unfold^42,50^. Overall, the intra-molecular *Gō*_intra_ captures the structural fluctuations observed in atomistic molecular dynamics as tracked by RMSF (Fig. S3). To assess how the differences in the description of the conformational flexibility of the folded domains impacts phase behavior, we run simulations in slab geometries as before for the EN model at *λ* = 1.05, *λ* = 1.06 and *λ* = 1.07 (Fig. S4). With the *Gō*_intra_ condensates are more stable at *λ* = 1.07 than with the elastic network model (Fig. 4A, Fig. S17 and Fig. S18). Across all conditions, the *Gō*_intra_ model consistently exhibits more pronounced phase separation than the EN baseline. This enhanced stability likely arises from increased inter-chain contacts captured by the *Gō*_intra_ model (Fig S16B and C), which strengthen the internal interaction network of the dense phase. We conclude that conformational flexibility can enhance phase separation^64^, possibly by enabling new inter chain interactions as proteins adapt their structure to the condensate environment, rather than disfavoring phase separation.

**FIG. 4.**
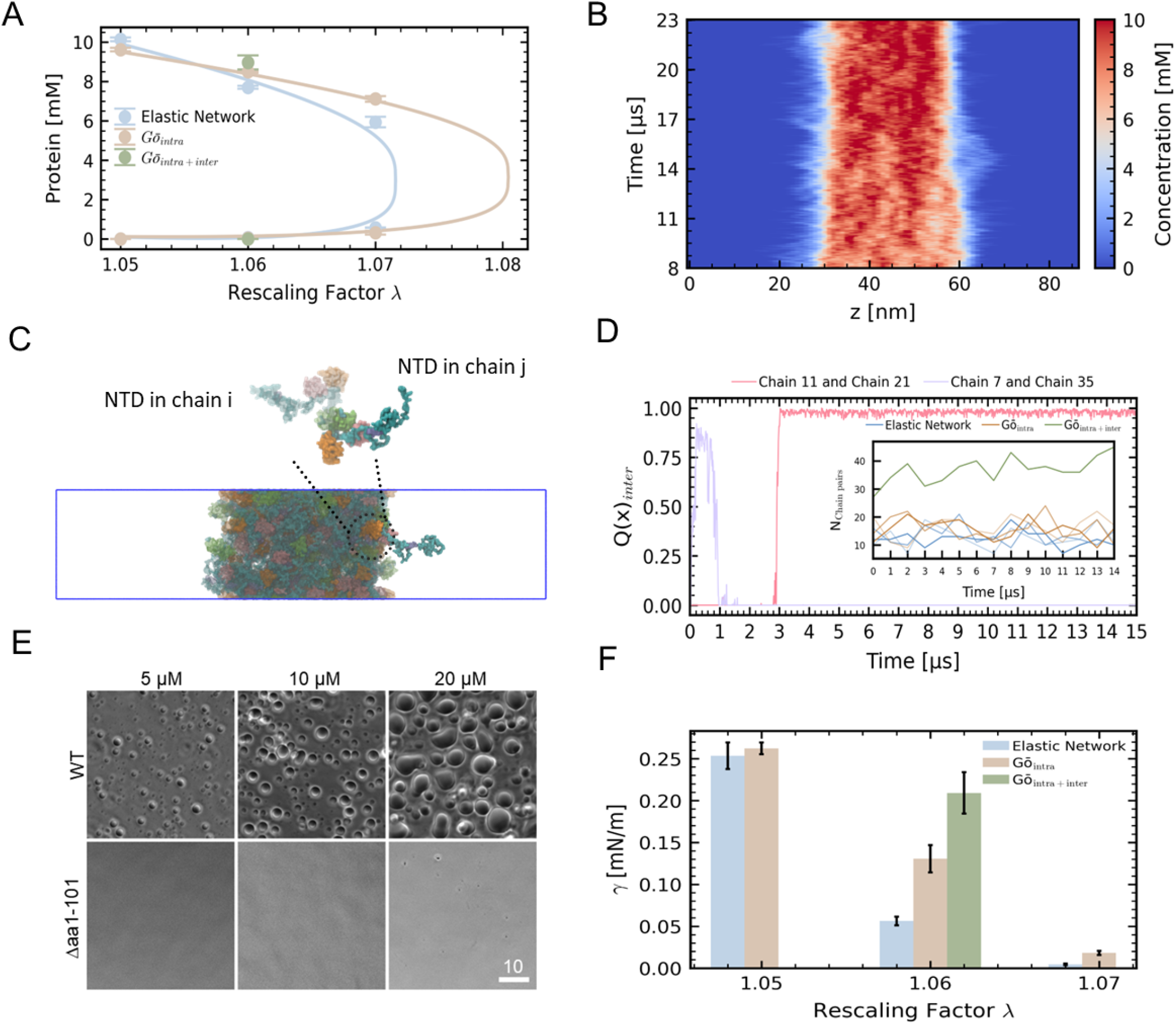
NTD-mediated intermolecular dimerization promotes TDP-43 phase separation and alters condensate material properties. (A) Phase diagram of EN, *Gō*_intra_ models along with dilute and dense phase concentrations at *λ* = 1.06 of *Gō*_inter + intra_ model. (B) Time-dependent density profile a Martini3 *Gō*_inter + intra_ model with protein-water interaction strength *λ* = 1.06 along the z-axis. (C) Representative condensate with NTD-dimer formed in the slab simulated by *Gō*_inter + intra_ model. (D) Inter-molecular *Q*(*x*) of the chain pairs within the condensate which form NTD-dimers, the insert presents the NTD-mediated chain pair number in three models. (E) *In vitro* experiments testing phase separation of full-length TDP-43 and a TDP-43 variant lacking residues 1–101 at different protein concentrations. (F) Surface tension measurement of EN and *Gō*_intra_ models across a range of *λ* values, alongside the surface tension of *Gō*_inter + intra_ model at *λ* = 1.06. The errors are propagated from the standard errors on the fitting parameters.

### Specific NTD-NTD interactions

Capturing highly-specific interactions of dimers and multimers, which depend on atomic-scale details is challenging in coarse-grained simulations, as key molecular details of the interaction interface are lost in the coarse-grained description. To establish a baseline for the dynamics of the NTD dimer, we performed atomistic molecular dynamics simulations on isolated NTD dimers. These simulations indicate that the crystallographically resolved NTD–NTD dimer interface is stable (Fig. S19B). However, in coarse-grained simulations of isolated dimers with both the EN and the *Gō*_intra_ models, although the specific interaction interface for NTD-dimerization is accessed, it does so only transiently. The fact that the native dimer structures is transiently accessed illustrates that the Martini3 models can capture specific protein-protein interactions at least in part (Fig. S19B and Fig. S20, Supporting Information). Since we wish to understand the how specific interactions of the NTD shape the properties of full-length TDP-43 condensates, we added additional Lennard-Jones interactions for the residues in the NTD-NTD interaction interface, resulting in a *Gō*_intra+inter_ model. First, we established that the *Gō*_intra+inter_ model promotes the specific dimerization of the isolated NTD in dilute solution as expected (Fig. S19 and Fig. S20, Supporting Information). Subsequently, we established that condensates of 60 TDP-43 full-length chains are maintained in simulation with the *Gō*_intra+inter_ model over durations beyond 20 *µ*s (Fig. 4B). Due to the enhanced NTD-NTD interactions more NTD-NTD mediated contacts between the full-length TDP-43 chains are formed. Within the condensate (Fig. 4B,C), specific NTD–NTD-mediated dimers dynamically form and dissociate, as illustrated by two representative examples in Fig. 4D, which shows a dimer that was formed then dissociated (chains 7 and 35), and another which formed after 3 *µ*s and remained stable until at least 15 *µ*s (chains 11 and 21). Our simulations show that the formation of stereospecific NTD–NTD dimers promotes TDP-43 phase separation consistent with *in vitro* experiments and favors additional interactions of the NLS, RRM2 and the LCD (Fig S16). The additional NTD-NTD interactions appear to diminish molecular exchange between the dilute and dense phases (Fig 4B). On the 20 *µ*s time scale we only see one chain leaving the condensate before quickly re-entering it. Multiple exchange events are observed for the baseline model, which is in line with the importance of the specific interactions of the NTD for the phase separation propensity of TDP-43. At *λ* = 1.06, the *Gō*_intra+inter_ model further enhances dense-phase stability, with an increased dense phase concentration. Intermolecular contacts beyond the NTD are enhanced (Fig S16B and C), and this increase goes beyond the additional interactions seen for *Gō*_intra_ model. These additional interactions likely help to suppress chain escape into the dilute phase. As TDP-43 dimers are formed, RRM2 inter-chain interactions become more prominent. NLS-NLS interactions are also enhanced by the NTD dimerization that brings the NLS regions from two TDP-43 chains close to each other, which suggests an interaction profile of the NLS region beyond the NLS-LCD contacts we established in simulations with the baseline EN model of full-length TDP-43.

Considering this new high-resolution molecular picture where NTD-NTD contacts enable further interactions, it is important to relate these simulations back to experiments. Indeed, *in vitro* experiments showed that a truncated variant lacking the NTD and NLS regions (Δ1–101) phase separates much less readily than both TDP-43 full-length and LCD. At a protein concentrations of 20 *µ*M, where the LCD formed amorphous structures, no Δ1–101 droplets were observed (Fig. 4E), which suggest that this truncation affects the intermolecular interactions of TDP-43 as least as much if not more than truncating the full-length protein to just the LCD (Supporting Text and Fig. S15). Conversely, it shows that contact statistics from such simulations can identify or prioritize sequence elements for truncation experiments. Based on the NTD and NLS mediated interactions in the simulations of the *Gō*_intra+inter_ model one would expect that such a variant is phase-separation deficient. This expectation is consistent with the observations from our *in vitro* assays of the truncated variant lacking the NTD and NLS regions (Δ1–101) where we did not observe phase separation. In line with these findings, abolishing NTD dimerization by mutating key residues in the interaction interface leads to phase separation deficient TDP-43^65^. Taken together, our results support the notion that N-terminal regions such NTD and NLS, engage in key intermolecular contacts in TDP-43 condensates, which enhance TDP-43 condensation.

### TDP-43 condensates are dynamic and their material properties are shaped by NTD dimerization

Besides the increased NTD-NTD interactions, the TDP-43 chains in the condensates remain dynamic. To characterize the dynamics of TDP-43 within the condensate phase across all three simulation models, we computed the mean-squared displacement (MSD), of monomers center-of-mass in *x, y* plane averaged over three independent replicas, as shown in (Fig. S21). All models display a degree of sub-diffusive behavior, with a slope less than 1 (Fig S22). Such sub-diffusive behavior is expected for dense protein condensate, since chains are trapped by interactions with other TDP-43 chains and confined to the condensate. Among them, the *Gō*_intra_ model displays reduced mobility relative to the other two models, reflecting stronger intramolecular constraints within the condensate phase. In contrast, the EN model exhibits the highest mobility, consistent with its reduced contact probability shown earlier. At intermediate timescales (≈ 1*µ*s), all models cross over to normal diffusion, indicative of relaxation of interaction network constraints at sufficiently long times. In the crossover to normal diffusion, the EN model seems to lag behind the other two models as indicated by the slightly lower MSD at late times. As a second measure of dynamics in the condensate, we computed the end-to-end autocorrelation function shown in inset of (Fig.S21). Consistent with MSD, the EN model displays the most rapid relaxation, while the other two models relax relatively slowly. The end-to-end relaxation in condensate is much slower than for TDP-43 in dilute solution relax on the sub *µ*s timescale (Fig.S23). The enhanced specific NTD dimerization in the *Gō*_intra+inter_ model has only a minor effect on global chain relaxation.

Next, we determined the surface tension of TDP-43 condenates from all three simulation models. The surface tension of protein condensates reflects the net balance between cohesive protein–protein interactions and solvation effects at the condensate boundary. As expected, the surface tension decreased with increasing values of *λ* (Fig. 4F, Fig. S24, and Fig. S25), reflecting that stronger protein–water interactions weaken the cohesive forces stabilizing the dense phase. For a fixed solvent interaction strength (*λ* = 1.06), the different models yield quantitatively distinct values for surface tension. The EN model exhibits a lower surface tension relative to both *Gō*_intra_ and *Gō*_intra+inter_ models (Fig. 4F), indicating that weaker inter-chain interactions (Fig S16) allow greater interfacial fluctuations. The interfacial tension is increased by the specific NTD dimerization in the *Gō*_intra+inter_ model, with an interfacial tension *γ* about 0.21 nm/m, while in the *Gō*_intra_ model, the interfacial tension is about 0.13 mN/m. In simulations without specific interaction and with the simple EN model of folded domains *γ* is much smaller, close to 0.06 mN/m.

## DISCUSSION

With near-atomic resolution coarse-grained simulations with an explicit description of water and ions^40^, we reveal how homo and heterotypic interactions drive the phase separation of the multi-domain protein TDP-43. Our simulations of full-length TDP-43 revealed many interactions centered around the LCD but also other domains. Our simulations predict that full-length TDP-43 phase separates much more readily than the LCD on its own. We confirm this difference in line experiments and quantify *c*_dilute_ for full-length TDP-43 compared to isolated LCD. However, considering the much shorter length of the LCD, it is still highly prone to phase separate, which is in line with previous investigations^11,12,25^ and the contact statistics from our simulations.

Importantly, our simulations revealed how phosphomimetic mutations and mutations of aromatic residues reduce the TDP-43’s tendency to phase separate. Simulations show that the phosphomimetic 12D variant phase separates less well than WT (Fig. 2I), with more chains found in the dilute phase (Fig. 2F and Fig. S9). Notably, Na^+^ ions accumulate more within the condensate, potentially contributing to partial neutralization of negatively charged residues (Fig. S11C,D and Fig. S12). The phosphomimetic mutations also re-organize the inter-chain interactions of the LCD. Contacts between the C-terminal ends of the LCD are lost (Fig. 2D), and more water molecules bind at the phospho sites (Fig. S12D). Lost interactions are compensated at least in part by new interactions with positively charged Lys and Arg groups (Fig. S12A). However, sampling limitations are still limiting the strength of the conclusions, which we have addressed and assessed with a Bayesian model. This assessment indicates a 77% probability that 12D phase separates less well than WT. The helical structure is maintained in these simulations, while the effect of phosphomimetic mutations on the stability of the conserved C-terminal helix has not been addressed. Meanwhile, the loss of secondary structure of the conserved C-terminal helix with no changes to the sequence robustly leads to a reduction in TDP-43 phase separation, with a 99% probability in the Bayesian model. However, condensates are more densely packed than WT with the helix preserved. This suggests that TDP-43 condensates may pack more efficiently in the absence of a structured C-terminal helix. On the other hand, replacing six Phe and three Trp residues with Ala, without preservation of the C-terminal helix, produces only a modest additional effect when compared to the WT sequence with no helix preservation. The change upon mutation is mostly reflected in *c*_*dense*_, with no major change in phase separation propensity. Although our simulations are performed on the full-length protein, the observed trend is qualitatively consistent with previous atomistic molecular dynamics simulations and NMR experiments for the isolated LCD^11,12,25^, which showed that the destabilization of the helix reduces the phase separation propensity of the LCD^11,12,25^.

Simulating large-scale biomolecular condensates at near-atomic resolution is challenging for multidomain proteins^45,48^, as it requires accurately capturing protein chain conformational dynamics and the detailed interaction interfaces that govern specific dimeric and multimeric assemblies. Although coarsegrained models enable faster simulations at larger scales, they necessarily sacrifice atomic-level detail that is essential for resolving protein flexibility and specific binding interactions. Modeling structured protein domains with elastic networks does not allow structural rearrangements, whereas the *Gō*_intra_ approach enables such rearrangements by allowing contacts stabilized by Lennard-Jones (LJ) potentials to break. Martini3 simulations using both EN and *Gō*_intra_ approaches to preserve the folded regions can partially reproduce specific NTD–NTD interfaces, as resolved by X-ray crystallography^16,17^. While their stability is clearly underestimated, as evidenced by simulations of isolated NTD pairs, it is nonetheless encouraging that the specific NTD–NTD complex can be formed in the coarse-grained simulations with the Martini3 model. To capture the dimerization of the NTD, we employed a *Gō*_intra+inter_ model developed by Korshunova et al^53^. This approach was previously used to study amyloid fiber formation^53^ and is a scalable and versatile way to model multimerization, using experimental and AlphaFold structures of complexes as an input. While the additional interactions between the NTDs are weak, we nonetheless observe more NTD-NTD contacts in the condensate. In principle these interactions could be tuned to match experimental interaction affinities.

In our simulations we observe how conformational flexibility and specific interactions of NTD dimers affect phase behavior, material and dynamic properties of condensates. Compared to the EN model, the *Gō*_intra_ model enables more conformational malleability, promoting more inter-chain contacts. Conformational flexibility thus enhances condensation as previously shown by Farr et al^64^. The interfacial tension of condensates is also increased by approximately a factor of two compared to the baseline EN model — from close to 0.06 mN/m to about 0.13 mN/m. Dimerization of the NTD within condensates further enhances NTD-NTD chain-to-chain interactions, increasing the interfacial tension of TDP-43 condensates from approximately 0.13 mN/m to 0.21 mN/m. The three models show similar diffusion behavior. While protein chains remain largely confined within the condensates on the microsecond timescale, the dynamics is sub-diffusive up the microsecond timescale, before transitioning into normal diffusion after that. The sub-diffusive behavior we have detected may limit the speed of enzymatic reactions in TDP-43 condensates as a protein substrate might take a long time time to encounter an enzyme in the condensate^66^.

Removal of the N-terminal residues (amino acids 1-101, encompassing both NTD and NLS)drastically reduces TDP-43 phase separation in the *in vitro* experiments (Fig 4E). Homotypic dimerization of the folded NTD is known to drive TDP-43 condensation, and the disruption of this interaction through mutations strongly reduces the phase separation propensity of TDP-43^65^. Loss of TDP-43 condensation is linked to defects in alternative RNA processing^9^. It has been proposed that loss of dimerization favors aggregation over condensation^17,20,65^, in line with the idea that these two pathways may be in competition with each other, as recently shown for the the LCD of hnRNP-A1^67^. In addition, one can speculate that loss of TDP-43 dimerization and hence condensation may lead to altered RNA-binding and RNA processing^9,62^. Our simulations reveal strong NLS–LCD inter-chain interactions, consistent with previous residue-level coarse-grained simulations ^39^, and in line with recent findings from experiments^68^, where NLS mutants reduced TDP-43 phase separation. Moreover, enhanced NTD–NTD interchain interactions also lead to increased NLS–NLS, RRM2–RRM2, RRM2–LCD and LCD–LCD interactions. The combined experimental and simulation results demonstrate that both the NTD and NLS are critical for promoting the multivalent interactions driving TDP-43 phase separation.

## METHODS

### Protein purification

His-Tev protease, full-length TDP-43-tev-MBP-His WT, W3A, F6W3A, as well as TDP-43 with an N-terminal deletion (Δaa 1–101), were purified according to Ref^69^ and Ref^8^, with the small modification that the storage buffer contained 2 mM TCEP instead of DTT. The TDP-43 LCD (aa 267–414)-tev-MBP was purified as a His-FKBP-3C-TDP43 (aa 267–414)-tev-MBP fusion protein^70^ (kind gift from Helle Ulrich, IMB Mainz) by the IMB protein production facility, and the His-FKBP tag was cleaved off during the purification process.

### Phase separation assays

Prior to all phase separation experiments, recombinant proteins were centrifuged for 10 min at 20,000 *g* and 4 °C to remove preformed aggregates. Phase separation assays were performed in phase separation buffer (20 mM HEPES, pH 7.5, 150 mM NaCl, 2 mM DTT). Condensate formation was analyzed approximately 1 h after addition of His-Tev protease at a fixed molar ratio (2.5× molar excess of TDP-43) and examined by phase-contrast microscopy using Ibidi *µ*-Slide 18-well flat chambers (ibidi, #81826) and a Zeiss Observer microscope.

To determine the concentration in the dilute phase upon condensate formation (*C*_sat_) for TDP-43 FL and LCD, condensate formation of a 10 *µ*M solution of the respective protein was induced as described above. After 60 min, samples were centrifuged for 10 min at 20,000 *g* and 4 °C. Equal fractions of pellet and supernatant were subsequently analyzed by SDS–PAGE and SYPRO Ruby total protein staining according to the manufacturer’s instructions. Band intensities of supernatant and pellet fractions were quantified by densitometry measurements (Image Lab software; Bio-Rad Laboratories). The relative amount of soluble protein was calculated as

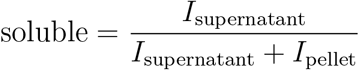

and the respective *C*_sat_ was assigned based on the known starting protein concentration (10 *µ*M) used in this assay^62^.

### Atomistic molecular dynamics simulations

Atomistic molecular dynamics simulations were performed using the GROMACS 2024^71–77^ software package with the AMBER99SB*-ILDN-q^78–81^ protein force field and the TIP4P-D^82^ water model. All systems were neutralized with physiological salt concentration of 150 mM NaCl. Periodic boundary conditions (PBC) were applied in all three spatial dimensions. Nonbonded interactions were computed using a 12 Å cutoff for both electrostatic and van der Waals (VDW) interactions, and long-range electrostatics were treated with the Particle Mesh Ewald (PME) method.

Equilibration was carried out in the NPT ensemble using a 2 fs timestep, allowing the system volume to relax before production simulations. During equilibration, temperature was maintained at 300 K using the Berendsen thermostat^83^, and pressure was regulated at 1 bar with the Berendsen barostat^83^.

Production simulations were subsequently performed in the NPT ensemble with a 2 fs integration timestep. Temperature and pressure maintained at 300K and 1 bar with the Bussi-Donadio-Parrinello velocity-rescaling thermostat^84^ and Parrinello-Rahman barostat^85^ respectively.

### Coarse-grained molecular dynamics simulations

Explicit solvent coarse-grained simulations were performed using the Martini 3 force field, or an adapted variant with rescaled protein–water interaction strengths^35,45^, within the GROMACS 2024 software frame-work. All Martini-based coarse-grained systems were solvated in water and neutralized with 150 mM NaCl. System equilibration was conducted under NPT conditions using a 10 fs timestep, with temperature maintained at 300 K via the Berendsen thermostat and pressure controlled at 1 bar with the Berendsen barostat.

Production simulations were performed using a 20 fs timestep. Temperature and pressure during production were maintained at 300 K and 1 bar, respectively, using the same velocity-rescaling thermostat and Parrinello–Rahman barostat as atomistic simulations. Box dimensions and pressure in XX and Z dimension were monitored during the simulations (Fig. S26, Fig. S27, Fig. S28).

#### Elastic network model

Multi-domain proteins contain both structured regions and intrinsically disordered regions (IDRs). The structured domains were stabilized using an Elastic Network (EN) Model, in which harmonic bonds are applied between coarse-grained beads to preserve the native fold.

Protein inputs (all-atom level) were coarse-grained by applying martinize2 in vermouth 0.14.1^86^ Python script. Secondary structures within the folded regions were assigned with the the STRIDE algorithm^87^ in the STRIDE Web interface. Elastic Network restraints were applied using a force constant of 700 kJ mol^−1^ nm^−2^, with bead pair distance included between 0 nm and 0.85 nm. The Elastic restraints (rubber bands), dihedral and angle potential within linker regions and IDRs between structured domains were all removed^35^, the structured regions applied with restraints are, residue 4 to 76, 105 to 176, 192 to 259 and 320 to 332.

The adapted Martini 3 force field was obtained by rescaling the protein–water interaction strength^35,45^ through modification of the Lennard–Jones (LJ) well depth, *ε*. Specifically, the LJ *ε* parameter between all protein and water beads were scaled by a factor *λ*. The Lennard–Jones potential is given by

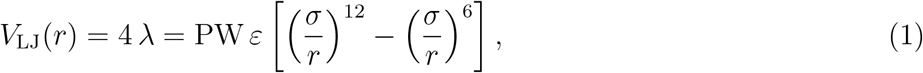

where *ε* is the interaction well depth, *σ* is the distance at which the LJ potential equals zero, and *λ* = PW is the scaling factor applied to protein–water interactions. The Python script used for protein– water interaction rescaling is available online at https://github.com/KULL-Centre/papers/tree/main/2021/Martini-Thomasen-et-al/PW_rescaling_example.

#### Intra Gō-type potential model

Following the Gō-like approach^50^, we stabilized native contacts in structured domains by creating virtual backbone beads and applying additional Lennard-Jones (LJ) potentials between relevant bead pairs. This section outlines the specific methodology for implementing LJ potentials for intra-chain contacts only within structured domains. The residue range definitions in the structured domains are the same as in the EN model.

Protein input structures were coarse-grained using martinize2 as described in the Elastic Network Model section. For simulation systems containing multiple chains, each chain requires a unique identifier for distinguishing. To define intra-chain native contacts, single-chain contact maps were obtained from a web server http://pomalab.ippt.pan.pl/GoContactMap/. The Python script create goVirt.py from https://github.com/Martini-Force-Field-Initiative/GoMartini/blob/main/GoMartini/create_goVirt.py was used to implement Gō-like bonds. The strength of the Go model structural bias was set to the default value of 9.414 kJ/mol. Additional statements were added to the Martini 3 force field to incorporate the created Gō-bonds. Similar to the Elastic Network Model approach, Gō-bonds, dihedral, and angle potentials within linker regions and intrinsically disordered regions (IDRs) between structured domains were removed. For multi-chain systems (e.g., 60 chains), the .tpr file generated in GROMACS for equilibration and production requires 128 GB of memory allocation.

Application of the adapted Martini 3 force field follows the same procedure as for the Elastic Network Model.

#### Intra and inter Gō-type potential model

For multi-chain systems, inter-chain contacts can be stabilized with additional Lennard-Jones (LJ) potentials alongside the intra-Gō potential. We adapted the approach by Korshunova and Bruiniks^53^for multi-domain proteins involving intrinsically disordered proteins (IDPs).

Similar to both the Elastic Network and intra Gō-type models, atom-level protein structures were firstly coarse-grained by using martinize2. The additional inter- and intra-chain LJ potentials were implemented using the adapted Python script `sorted goVirt.py` from the same repository. This script can exclude LJ potentials for linker regions and IDRs upon request. Additionally, dihedral and angle potentials within these flexible regions between structured domains have to manually remove.

This approach can also be utilized for systems requiring only the Intra Gō-type potential model, which offers the advantage of reduced memory allocation compared to multi-chain implementations.

#### Multi-chain simulation setup

Multiple full-length TDP-43 chains were simulated in both cubic and slab geometries. 70 Protein chains were initially placed at random positions within a cubic simulation box (50*nm* × 50*nm* × 50*nm*, which was subsequently solvated with water and neutralizing ions. Systems in the cubic geometry were simulated using the adapted Martini 3 force field with a rescaling factor of *λ* = 1.06 during energy minimization, equilibration, and production runs.

Simulations in slab geometry were run for 60 chains per simulation system. Simulation in the slab geometry were started from different configurations, the initial configurations were listed in the Fig S4. These starting structures were generated from preliminary condensation runs using an adapted Martini 3 force field. The simulation box was initially set to 20 nm × 20 nm × 90 nm for equilibration, ultimately stabilizing at 86.7 nm × 19.7 nm × 19.7 nm.

#### Simulations in dilute solution

Simulations of single TDP-43 chains were performed using both atomistic and Martini 3 coarse-grained models. The Martini 3 simulations were carried out at *λ* = 1.00 and *λ* = 1.06. For each condition, three independent replicas were run, each exceeding 10 *µ*s in duration.

### Analysis of phase coexistence in coarse-grained molecular dynamics simulations

The quantitative analysis of phase behavior was performed on simulations employing a slab geometry. Density profiles were determined from the Martini3 simulations following the work of Tesei et al^58^.

The system was aligned by translating the protein chains such that their collective center of mass (COM) was centered at *z* = 0. The spatial distribution of protein along the z-axis was computed and stored as a histogram for each trajectory frame. The time-averaged density profile was then symmetrized by averaging the density from both slab halves (*z <* 0 and *z >* 0), producing a single symmetrized profile for *z* ≥ 0. This symmetrized profile was fit to a hyperbolic tangent function to characterize the interface:

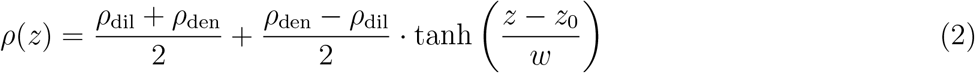

where *z*_0_ defines the midpoint of the interface and *w* parameterizes the interface width. The coexisting phase concentrations, *ρ*_dil_ and *ρ*_den_, were determined directly from the averaged histogram by calculating the mean density in regions defined by the interface cutoffs. To estimate statistical uncertainties, the trajectory was divided into segments. The average concentration within each block was computed, and the standard error of the mean (SEM) was calculated from these block-averaged values.

### Excess Free Energy of Transfer from the Dilute to the Dense Phase

To quantify how favorable it is for one chain to move from the dilute phase to the dense phase at the saturation concentration, we compute the excess free energy of transfer

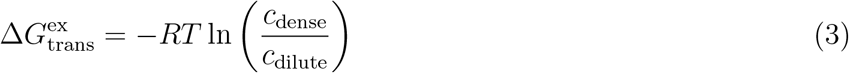

^47^ (Supporting Information). A more negative value of 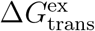 indicates a higher phase separation propensity, whereas a more positive 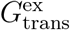 indicates that is less favorable to move one chain from the dilute phase to the dense phase.

### Bayesian Estimates of the Excess Free Energy of Transfer

To assess the relative propensity of TDP-43 variants to phase separate we calculate 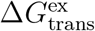 based on the mean concentration *µ*_*c*_ in the dilute and dense phase. To estimate *µ*_*c*_ for a single variant we need account for systemic qualitative differences in individual replicate simulation runs (e.g. durations or sampling speed). We do this by using a Bayesian approach, that incorporates both the individual standard error *ϵ* of a single simulation as well as a shared nuisance parameter *τ* across replicate simulation (pooling).

In the following *j* denotes the variant and *i* the (replicate) simulation index. For each simulation *i* we analyze the dilute (dil) and dense (den) phase as log concentrations log_dense*/*dilute,*i*_ sampled from slab density time series *c*_*i*_(*z, t*).

The individual uncertainty *ϵ*_dense*/*dilute,*i*_ was estimated by *B* = 10 000 bootstrap resamples of *N*_*i*_ contiguous equal-length blocks, with one bootstrap for each of the two phases.

We define (fixed) Gaussian priors for the log latent variant-level dilute and dense phase concentrations log *µ*_*c*,dil,*j*_ and log *µ*_*c*,den,*j*_,

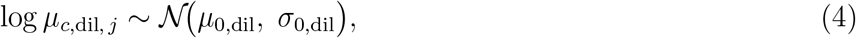

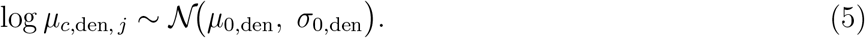

We set the hyperparameters *µ*_0,dilute_ = −2.3, *σ*_0,dilute_ = 2.0, *µ*_0,dense_ = +2.7, and *σ*_0,dense_ = 1.5 (standard deviations). The 95 % prior intervals of *µ*_*c*,dilute_ ∈ [2 × 10^−3^, 5] mM and *µ*_*c*,dense_ ∈ [0.8, 290] mM imply a range of 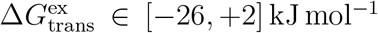 at the 1–99 % range, which spans a large range of physically plausible values.

We include nuisance parameters *τ*_log,dil_ and *τ*_log,den_ that capture inter-simulation scatter beyond the per-simulation block-bootstrap uncertainty in the log dilute and dense concentrations *ε*_log,dilute,*i*_ and *ε*_log,dense,*i*_

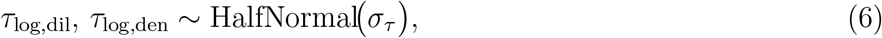

with the hyperparameter *σ*_*τ*_ = 0.5.

The total log-space uncertainty for each observation is

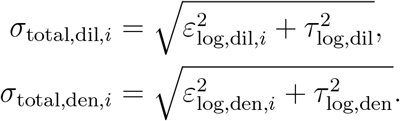

We define the likelihood for the log dilute and log dense concentrations as Normal distributions centered on the respective variants latent log concentrations log *µ*_*c*,dil, *j*_. Thus,

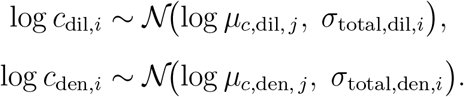

The excess free energy of transfer (Eq. 3) for each variant was then computed from the latent concentrations,

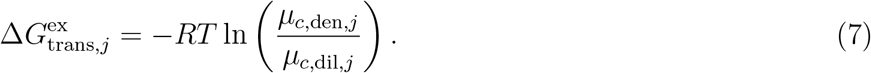

The change in phase-separation propensity of variant *j* relative to WT is then

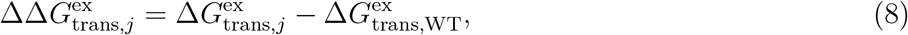

which we report with Bayesian credible intervals.

To quantify how strongly the simulations support the conclusion that a variant *j* phase-separates less well than the WT protein we compute the proportion of the posterior distribution where 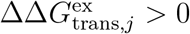,

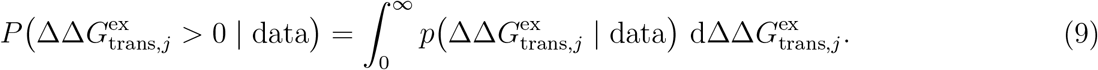

We used Markov Chain Monte Carlo in PyMC5, with the No-U-Turn Sampler for inference. Four chains were used, with 2000 independent samples and 2000 tuning steps. Convergence was assessed with the Gelman-Rubin statistic 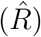, which was *<* 1.01 for all parameters of our model. The initial Bayesian approach was developed with the help of Google Gemini 2.5 Pro and refined with Claude Sonnet 4.7 and Opus 4.8

### Construction of phase diagrams

Phase diagrams were constructed by plotting the coexisting phase concentrations, *ρ*_dil_ and *ρ*_den_, against the interaction rescaling factor *λ*. The critical point (*λ* = *C*) was obtained by

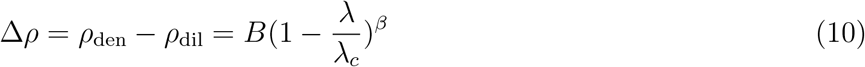

B is a fitting parameter representing the critical amplitude, and assume the systems belong to the 3d Ising universality class, with a critical exponent *β* = 0.325.

The critical density *ρ*_*c*_ was estimated using the law of the rectilinear diameter:

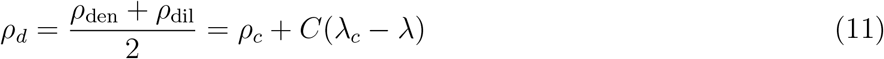

is applied with a positive fitting constant C^88,89^.

### Analysis of conformations in atomistic and coarse-grained molecular dynamics simulations

Native contacts for the inter-chain interactions were adapted and computed following Best and Hummer^90^ for atomistic and coarse-grained simulations of TDP-43 NTD dimer.

The geometric *R*_*G*_, *i* values for structures *i* sampled in simulations were computed from the single-chain simulations with the MDAnalysis Python library. The average^91^ *R*_*G*_ was determined by:

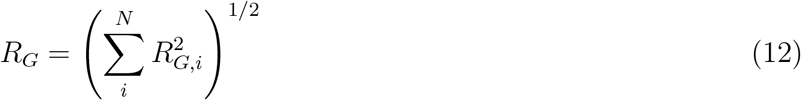

The SEM was estimated using block analysis.

### Interfacial tension calculations

Estimation of condensate surface tension is based on Capillary wave theory^92^ (see SI for detailed description) with block analysis method in the slab geometry^93^. The idea is to relate the surface tension to fluctuations in the interfacial width of the condensate. To do so, we divide our system into blocks of length *B* in the lateral (x-y) direction. Each block spans the entire box in the z-direction. Each block then has dimensions *B* × *B* × *L*_*z*_. In each block, we calculate the density profile and fit it to

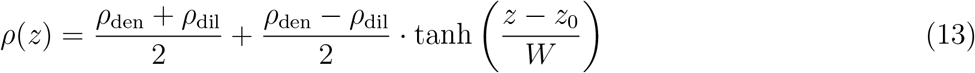

where *ρ*_den_ is the density in the dense phase, *ρ*_dil_ is the density in the dilute phase, *z*_0_ is the position of the interface, and *W* is a parameter related to the width of the interface. Following arguments based on capillary wave theory, we can show the relation^92,93^ (see also SI)

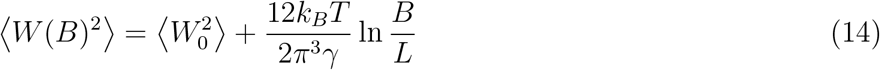

where *W* (*B*) is the fitted width parameter for blocks of length *B, W*_0_ is the width parameter for *B* = *L, k*_*B*_ is the Boltzmann constant, *T* is the temperature, and *γ* is the surface tension. The surface tension can then be calculated by measuring ⟨*W* (*B*)^2^⟩ for different block sizes *B*. In practice we choose block sizes 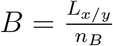 with the number of blocks *n*_*B*_ ∈ [4, 10]. The results are shown in Fig. S24 and Fig. S25.

### Mean-squared displacement

The lateral dynamics were quantified using the ensemble-averaged mean-squared displacement (MSD) in the plane parallel to the slab interface

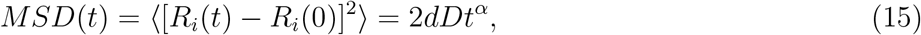

where ⟨..⟩ indicates the ensemble average, *R*_*i*_ is the backbone (BB) coordinates. The diffusion coefficient is given by *D* and *d* = 2 indicates the dimensions of the system. For normal diffusion in the lateral plane, we expect MSD to grow linearly with *α* = 1 in the long time limit. For the efficient computation of the MSD we used a blocking scheme^94–96^ which is a modified version of the “order-n-algorithm^96^.

## ACKNOWLEDGMENTS

This project was funded by the Deutsche Forschungsgemeinschaft (DFG, German Research Foundation) - SFB 1551 – Project No. 464588647. X.P. acknowledges a PhD fellowship from the Max Planck Graduate Center (MPGC) Mainz. X.P. was funded by the Deutsche Forschungsgemeinschaft (DFG, German Research Foundation), Project No. 233630050 – TRR 146. L.S.S. and X.P. thank M^3^ODEL for support. L.S.S gratefully acknowledges support from ReALity (Resilience, Adaptation and Longevity) and Forschungsinitiative des Landes Rheinland-Pfalz. We thank Sebastian Thallmair, Paulo Telles de Souza, Yannick Witzky, Adolfo Poma, Martin Girard, Camilo Aponte-Santamaría, Oleksandra Kukarkenko, Kumar Gaurav, Mahesh Yadav, Emanule Zippo, Arya Changiarath, and Supriyo Naskar for insightful discussions. We acknowledge the SFB1551 internal service project Z01 (Biopolymer Engineering and Bioanalytics at IMB’s Protein Production Core Facility). We are grateful to Philipp Schönberger and Helle Ulrich (IMB Mainz) for providing the plasmid encoding His-FKBP-TDP-43 LCD (aa 267–414)-tev-MBP.

## DATA AVAILABILITY

The trajectories used for the analyses are available on Zenodo(https://zenodo.org/records/20597321).

## CODE AVAILABILITY

The Martini 3 Elastic Network and Gō-intra+inter model setup files, together with the code used for the Bayesian assessment of the phase-separation propensities of TDP-43 variants, are available on Zenodo (https://zenodo.org/records/20672935).

## SUPPORTING INFORMATION

### SUPPORTING TEXT

#### Definition of the Excess Free Energy of Transfer

At equilibrium the chemical potential of the dense and dilute phase match *µ*_dense_ = *µ*_dilute_. We split the chemical potential into two terms, *µ*^id^, an ideal gas term that depends only on the concentration, and *µ*^excess^, which includes contributions from interactions.

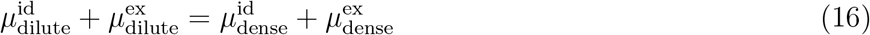

Rearranging gives

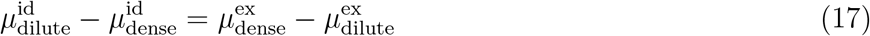

The non-ideal contribution is the excess transfer free energy, defined as

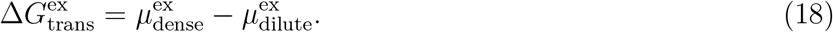

The excess transfer free energy 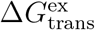 is a measure of the contribution of interactions to the phase separation. Since the ideal chemical potential is given by *µ*^id^(*c, T*) = *RT* ln *c* + *C*(*T*) where *c* is the concentration and *C*(*T*) is a temperature-dependent constant, we obtain at equilibrium

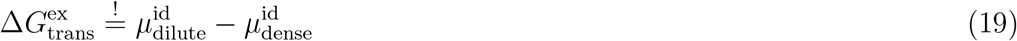

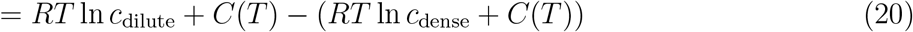

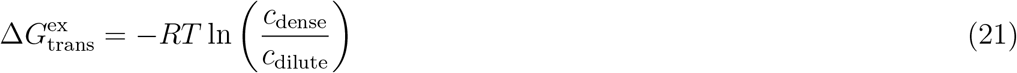

#### Capillary wave theory for flat interfaces

In a two phase system with a flat interface separating the phases, two useful parameters related to the interface are its position *z*_0_ and width Δ_0_. For a system at non-zero temperature *T*, the interface will be subject to thermal fluctuations that induce capillary waves. As a result, the interface will on average appear to be broadened with a width Δ. Assuming that the fluctuations only affect the position of the interface *z*_0_ but none of the other intrinsic properties, the fluctuation in the position of the interface can be found as (Refs. 97 §4, 98 Eqs. 8,9, and 92 Eq. 4)

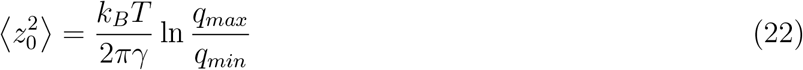

where *γ* is the surface tension, and *q*_*min*_ and *q*_*max*_ are upper and lower cutoff values when evaluating the integral over oscillation modes. *q*_*min*_ is related to the size of the system, while *q*_*max*_ is related to a molecular length scale. This fluctuation of *z*_0_ will cause the broadening of the interfacial width. If the order parameter in the absence of thermal fluctuations can be described through a profile *ψ*_0_(*z*), the broadened profile Ψ(*z*) can be obtained as the average of the intrinsic profiles *ψ*_0_(*z*) with different positions *z*_0_ of the interface, weighted by the probability *P*(*z*_0_) of the interface being at position *z*_0_

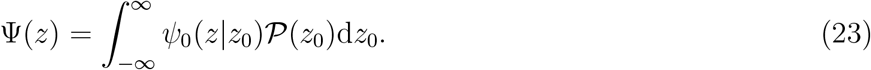

Lacasse et al. derive a relation between the broadened width Δ, intrinsic width Δ_0_, and the fluctuation in the position of the interface 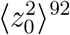

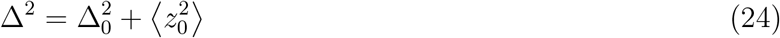

where the widths are defined through the measure^92^ (Eq. 6)

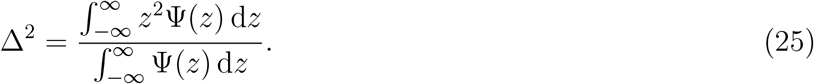

Replacing Eq. 22 in Eq. 24 and setting *q*_*min*_ = 1*/L* and *q*_*max*_ = 1*/B*_0_ we get

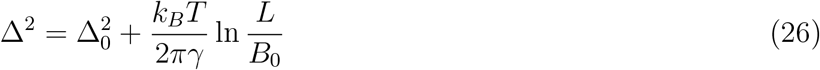

where *L* is the macroscopic length of the system and *B*_0_ is an undetermined molecular length scale. Now the remaining step to implement this is to find the width Δ from simulation. This depends on the choice to represent the density profile as described below.

##### The hyperbolic tangent profile

Following mean field theory considerations^99^, it can be found that the density profile *ρ*_0_(*z*) of a liquid phase with density *ρ*_*h*_ coexisting with a vapor phase with density *ρ*_*l*_ takes the form

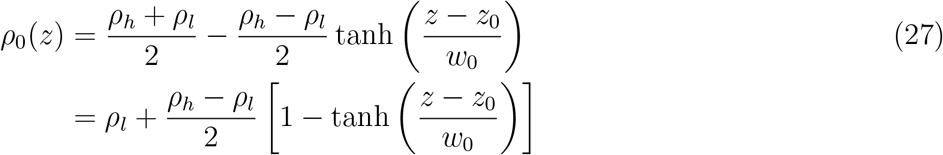

where *z*_0_ is the position of the interface and *w*_0_ is a parameter related to the interfacial width. An order parameter can be defined as

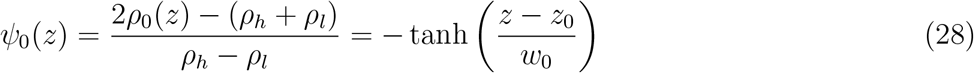

which takes values between −1 for the dilute phase and 1 for the dense phase. The broadened density profile is still well approximated by the same functional form but with a new width parameter *w > w*_0_. Using Eq. 28 in Eq. 25 we get for the intrinsic and broadened widths^92^

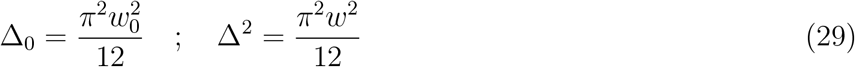

which gives the relation

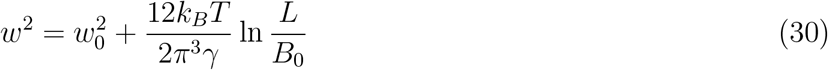

##### The error function profile

For completeness, we also give the result for the error function profile sometimes used in the literature^47,92,100^. The density profile is taken to be of the form

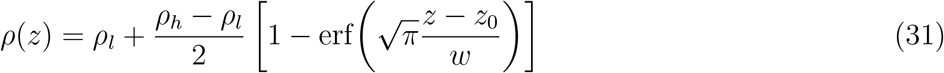

in which case the relations becomes^92^

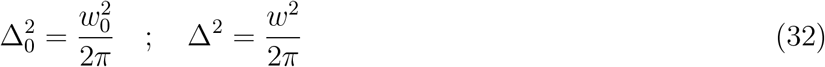

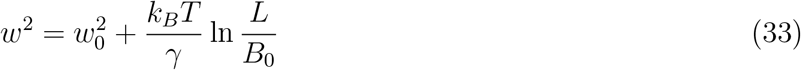

Note that in Refs. 47,100 the numerical factor in the erf() is taken as 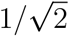 instead of 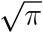

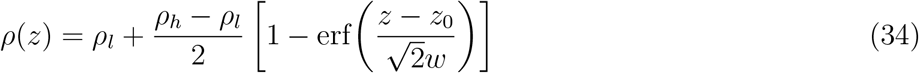

which leads to

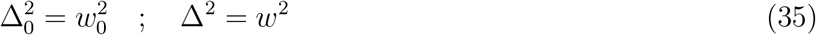

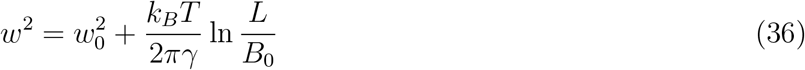

#### Global dimension of TDP-43 in dilute solution

Repeated simulations using the Martini 3 EN and *Gō*_intra_ models show consistent behavior in the radius of gyration (*R*_*G*_) for single-chain full-length TDP-43 (Fig S1). Simulations with both simulation models at *λ* = 1.06 give *R*_*G*_ values much closer to experimentally determined *R*_*G*_ than all atom simulations. The all-atom simulations transiently access more extended structures with *R*_*G*_ values close to experiment. Longer simulation durations would be required to assess the deviation from experiments due to remaining in balances in the force field^91^.

Example structures from these simulations all-atom simulations, EN and *Gō*_intra_ simulations are shown in Fig. S2. These structures are close to the average *R*_*G*_ in the simulations. As such they give view of the global conformation of TDP-43 in these simulations, but it is important to note that for multi-domain protein such as TDP-43, with extended IDR regions, a single structure is by necessity an incomplete representation of the structural ensemble.

#### Secondary structure of TDP-43 in atomistic and coarse-grained simulation in dilute solution

Residue flexibility of the folded domains of TDP-43 as seen in atomistic molecular dynamics is captured by simulations with the rescaled Martini3 model and an elastic network (EN) to capture the dynamics of folded domains. This flexibility is quantified by the root-mean-square fluctuation (RMSF) in the positions of the backbone beads in coarse-grained models or of the C*α* atoms from each residue in all-atom simulations. Atomistic molecular dynamics simulations in general provide a very accurate description of the conformational ensembles of folded protein domains^50,101^ and thus can serve as a benchmark for Martini3 models^50,101^. Atomistic molecular dynamics simulations started from an AlphaFold structure show that the secondary structure of the folded NTD, RRM1, and RRM2 domains is stable over the *µ*s simulation time scale. The color at different time points represents whether the particular residue is part of a helical, an extended (*β*-sheet), or an unstructured sub-region. Residues at the beginning and the ends of extended (*β*-strands like structures) and helical structures repeatedly switch between regular secondary structure and coil like formations, for instance residues close to 136 and 143-146 in RRM1. Residues 318 to 329 are mostly in helical conformations in line with NMR experiments and previous all-atom molecular dynamics simulations^11,12,25^ (Fig. S3D). Residues 330 to 333 also adopt helical conformations, for instance starting at about 0.6 *µ*s, which persists for the remainder of the trajectory.

In Figure S3 the RMSF per residue is shown for the four folded regions in TDP-43, NTD (Fig. S3A), RRM1 (Fig. S3B), RRM2 ((Fig. S3C), and the conserved helix (Fig. S3D) in the C-terminal LCD. Overall the level of conformational flexibility from atomistic molecular dynamics is captured for all four regions, in particular for regions with regular secondary structure. For the NTD, the EN approach overestimates the flexibility of residues 45 to 49, which are in a coil region flanked by regular extended (*β*-sheet like structure). Residues maintaining stable secondary structures within the sub-folded region show low RMSF, while the loops maintain flexibility in the EN Martini3 simulations, resembling the behavior observed in atomic models.

For instance, residues 216-221 in RRM2 stabely form an extended *β*-strand like conformation in the atomistic molecular dynamics simulations and RMSF values are below 2 Å for these residues. By contrast, focusing on the *α*-helix formed by residues 236-242, we find that some residues towards its N- and C-terminal ends transiently lose their helical conformations; nevertheless, the RMSF values remain low as the segment keeps its structure. The N-terminal residues of the adjacent *β*-strand residues 246-250 sometimes lose their extended conformation and the RMSF values for these residues rise to values close to 2 Å.

Simulations with Gō-like models shows similar behavior as EN model for simulations of single TDP-43 chains.

#### Comparison of *c*_dilute_ from phase separation experiments

We plot the experimental dilute-phase concentration *c*_dilute_ for full-length TDP-43 and for the LCD construct, together with the lower-bound estimate for the *δ*101 construct (which did not phase-separate in the brightfield droplet assay up to 20 µM), against their residue number *N* (Fig. S15). For comparison we overlay how *c*_dilute_ would scale with *N* for a homopolymer at fixed Flory–Huggins interaction parameter *χ* ∈ 0.58, 0.60, 0.63, 0.650.58, 0.60, 0.63, 0.65, computed from the common-tangent construction on the Flory–Huggins free energy. Because the dilute arm scales roughly as *c*_dilute_(*N*) ∝ exp [−(*χ* − *χ*_*c*_(*N*))*N*] at large *N*, longer chains phase-separate at lower concentration at any fixed *χ*. A construct that phase-separates more strongly than expected from chain length alone falls below the homopolymer *χ*-line passing through any reference construct of larger *N*; one that phase-separates less strongly than expected falls above it. The LCD’s *c*_dilute_ = 4.5 *µ*M at *N* = 154 sits well below the *χ* = 0.63 line that passes through the full-length point at (*N, c*_dilute_) = (414, 0.51*µ*M), suggesting that on a per-residue basis the LCD is substantially more cohesive than the chain-averaged full-length TDP-43.

#### NTD dimerization

The N-terminal domain (NTD) of TDP-43 is capable of forming homodimers, predominantly mediated through interactions between the head region of one monomer and the tail region of a neighboring monomer^17^. The dimerization interface is characterized by electrostatic complementarity, where a negatively charged head region interacts with a positively charged tail region, facilitating stable association. Structural insights from the crystal structure (Figure S19A) show that the tail *β*-sheet (*β*_51−58_, colored pink) establishes contacts not only with the head *β*-sheet (*β*_16−20_, pink) but also with the *α*-helix (*α*_26−35_, green) of the adjacent chain.

The NTD homodimer was simulated using all-atom models, the EN model, and both coarse-grained *Gō*_intra_ and *Gō*_intra+inter_ models. The fraction of native contacts, *Q*_(*x*)_^90^, was used to quantify the preservation of native inter-molecular contacts within the NTD dimer. In all-atom simulations, the dimer remained stably associated over the full 2 *µ*s trajectory, with *Q*_*x*_ values consistently close to 1 (Figure S19B), in excellent agreement with the reported experimental crystal structure^17^ (PDB: 5mdi).

All three coarse-grained models were able to capture NTD dimer formation, though with varying degrees of accuracy in reproducing the native inter-chain contacts (Figure S19B and S20). The decomposition of native contacts focused on the specific interactions between the *β*_51−58_–*β*_16−20_ and *β*_51−58_–*α*_26−35_ interfaces, as well as other residue contacts across the dimer. The EN model successfully captured the overall formation of the dimer; however, it rarely reproduced both the *β*_51−58_–*β*_16−20_ and *β*_51−58_–*α*_26−35_ interfacial contacts with consistently high *Q*_(*x*)_ values. This indicates that, while the EN model is sufficient to maintain global dimer architecture, it may not fully capture the detailed residue-level interactions that stabilize the interface.

The *Gō*_intra_ model also maintained the dimer and sampled interfacial interactions more frequently than the EN model, suggesting improved flexibility in capturing transient contacts. Nevertheless, the two critical contact interfaces remained somewhat less stable than in the all-atom reference, highlighting limitations in describing inter-chain specificity when only intra-chain interactions are included. In contrast, the *Gō*_intra+inter_ model, which incorporates additional inter-chain Lennard–Jones potentials to better describe native contacts, consistently preserved dimer formation and produced relatively high *Q*_(*x*)_ values across both interfaces. Despite this improvement, the *β*_51−58_–*α*_26−35_ contacts were still somewhat underestimated, suggesting that even the enhanced coarse-grained model cannot fully recapitulate all atomistic details of the NTD dimer interface.

#### Box dimensions and pressure

We monitored box dimensions and pressure in the simulations of the slab systems. The simulations were started from configurations pre-equilibrated in the slab geometry; thus, we expect that the box dimensions will be close to their respective equilibrium values. For both the EN and *G*ō_intra_ models, we observe that the pressure along the *z*-axis fluctuates around 1 bar (Fig. S26). The simulations with the EN model at *λ* = 1.06 show only very small fluctuations in the length of the *z*-axis of the box (Fig. S27). Somewhat larger fluctuations are observed for *L*_*x*_ and *L*_*y*_, but these are still below 1 Å. The same behavior also holds for simulations with the *G*ō_intra_ model at *λ* = 1.06 (Fig. S28).

## I. SUPPORTING FIGURES

### SUPPORTING MOVIES

Movie S1: Simulation of TDP-43 in the dilute phase.

Movie S2: Phase separation of TDP-43 in a cubic simulation box.

Movie S3: Simulations of the TDP-43 condensate in slab geometry.

**FIG. S1.**
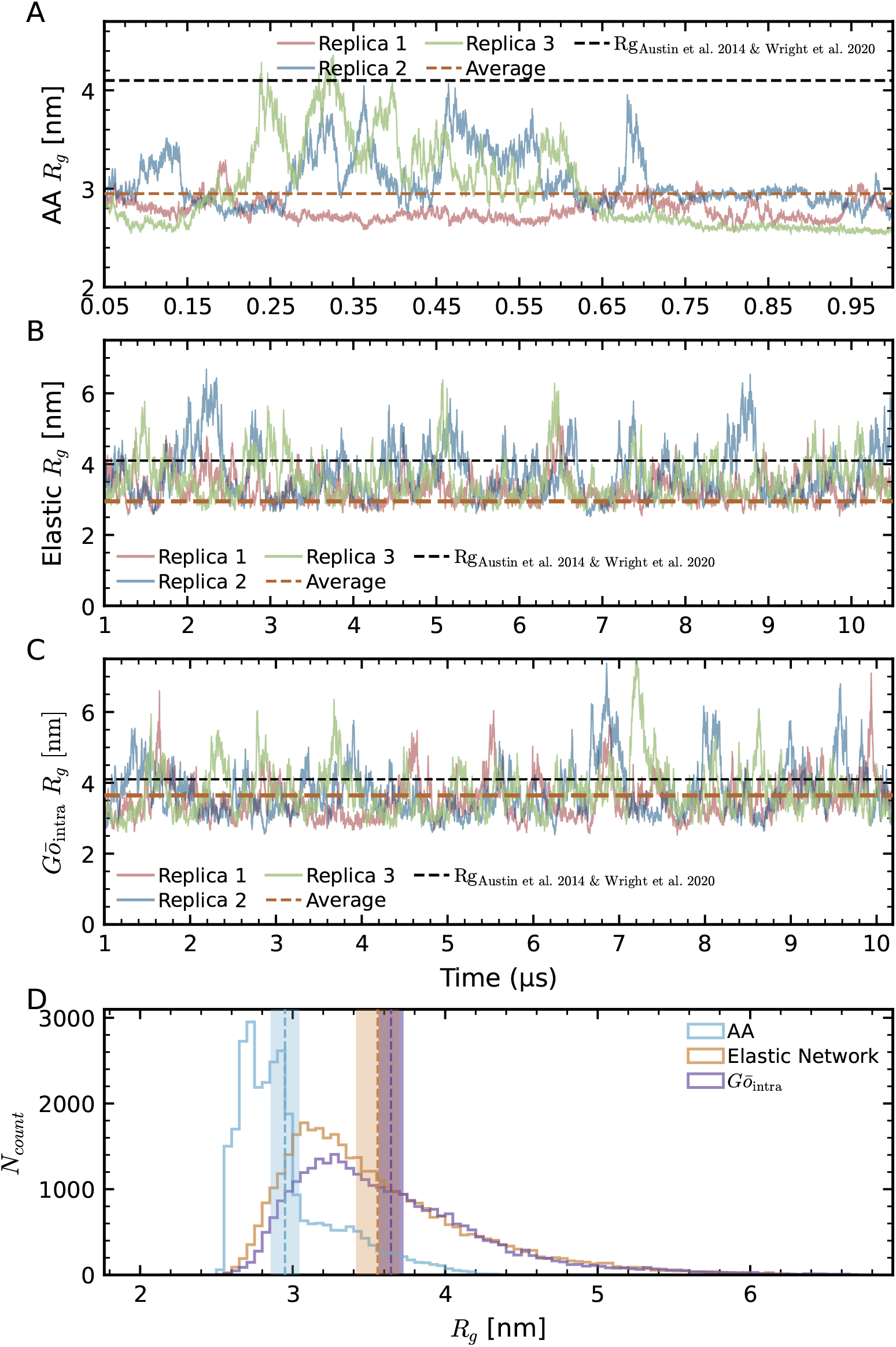
Characterization of full-length single-chain TDP-43 in all-atom (AA) and Martini3 coarse-grained (CG) simulations. The Martini3 CG model implements either an Elastic Network or *Gō*_intra_ model for the folded domains. (A–C) Time evolution of the TDP-43 radius of gyration (*R*_*G*_) from AA and Martini3 CG molecular dynamics (MD) simulations. CG models were simulated with strengthened protein-water interactions, scaled by factors of 1.06 (*λ* = 1.06). AA model results are shown for three independent replicas, each distinguished by different colors; brown dashed lines indicate the average *R*_*G*_ for each model. The black dashed line represents the maximum *R*_*G*_ value derived from experimental data. (D) Histograms showing the *R*_*G*_ distributions, calculated as the sum over all replicas for each simulation model.

**FIG. S2.**
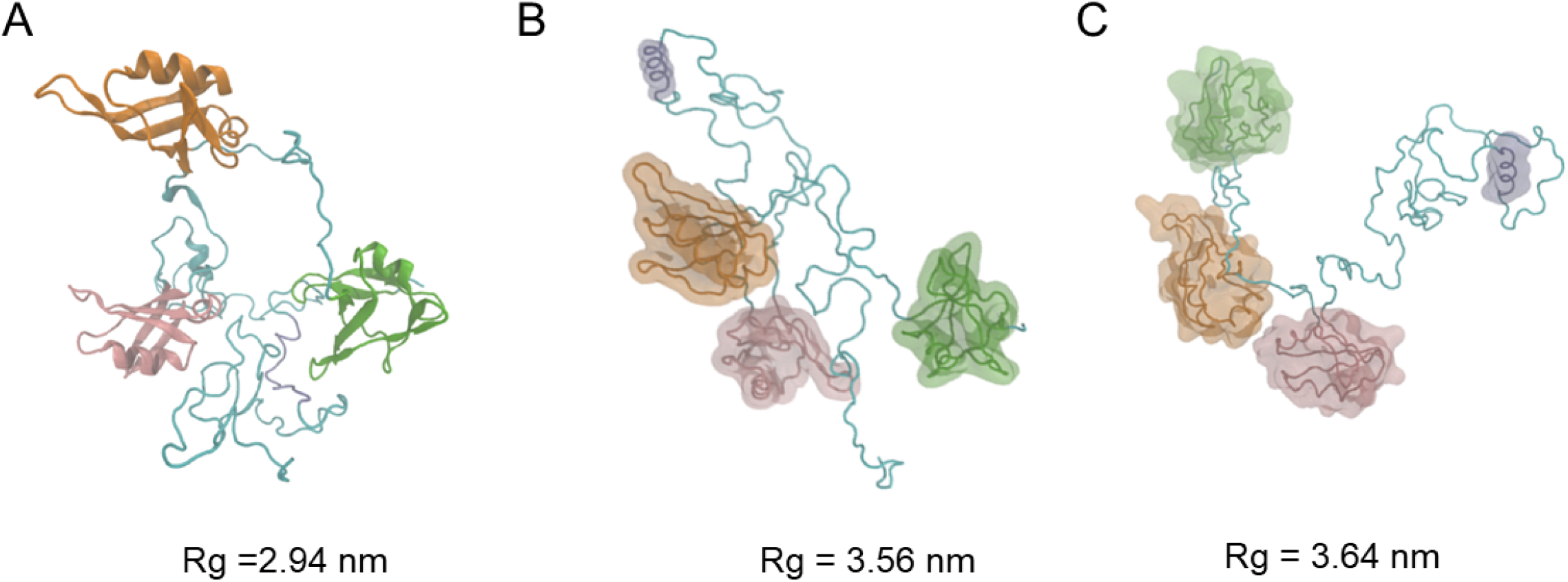
Representative structures from AA, Martini3 elastic network, and *Gō*_intra_ AA models, with average Rg highlighted. Residues highlighted in green, orange, pink correspond to the N-terminal domain (NTD), RNA-recognition motif 1 (RRM1), and 2 (RRM2), respectively. Cyan areas indicate the protein’s disordered regions (IDR), and the region colored in purple represents the helical segment within the C-terminal IDR. water and ions are omitted for clarity for both AA and Martini3 models

**FIG. S3.**
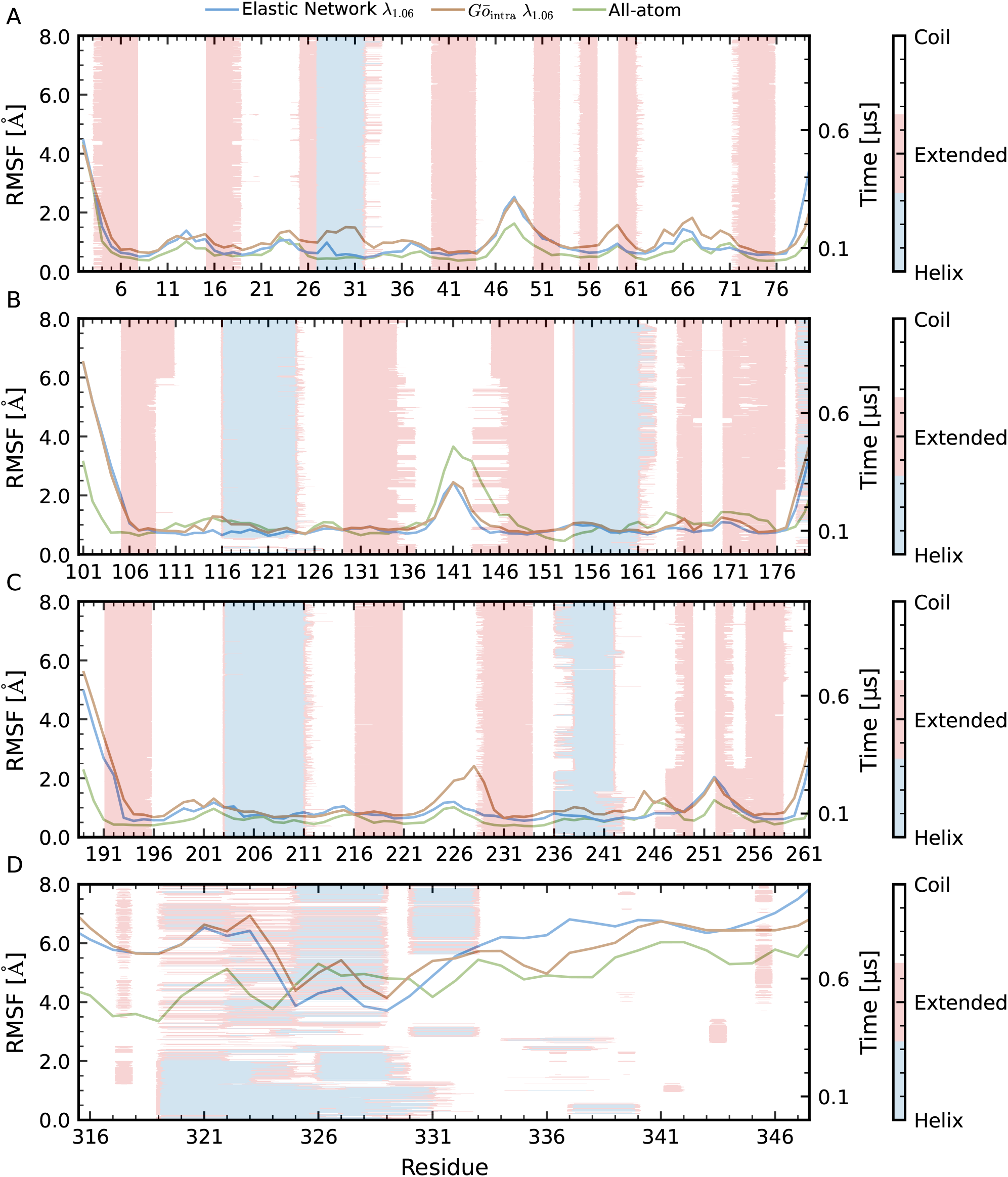
Residue flexibility within the structured domains of TDP-43 under all-atom, Elastic Network and *Gō*_intra_ models at *λ* = 1.06. Root mean square fluctuation (RMSF) of backbone beads in folded regions: NTD, RRM1, RRM2, and the helical segment of the C-terminal disordered region. The left axis measures the RMSF of each residue from its average position. Here both rescaled and unrescaled Martini 3 models are shown for comparison. The right axis in combination with the colorbar track the secondary structure of each residue over the time course of an AA simulation: blue is helix, pink is extended structure (*β*-sheet), and white is coil.

**FIG. S4.**
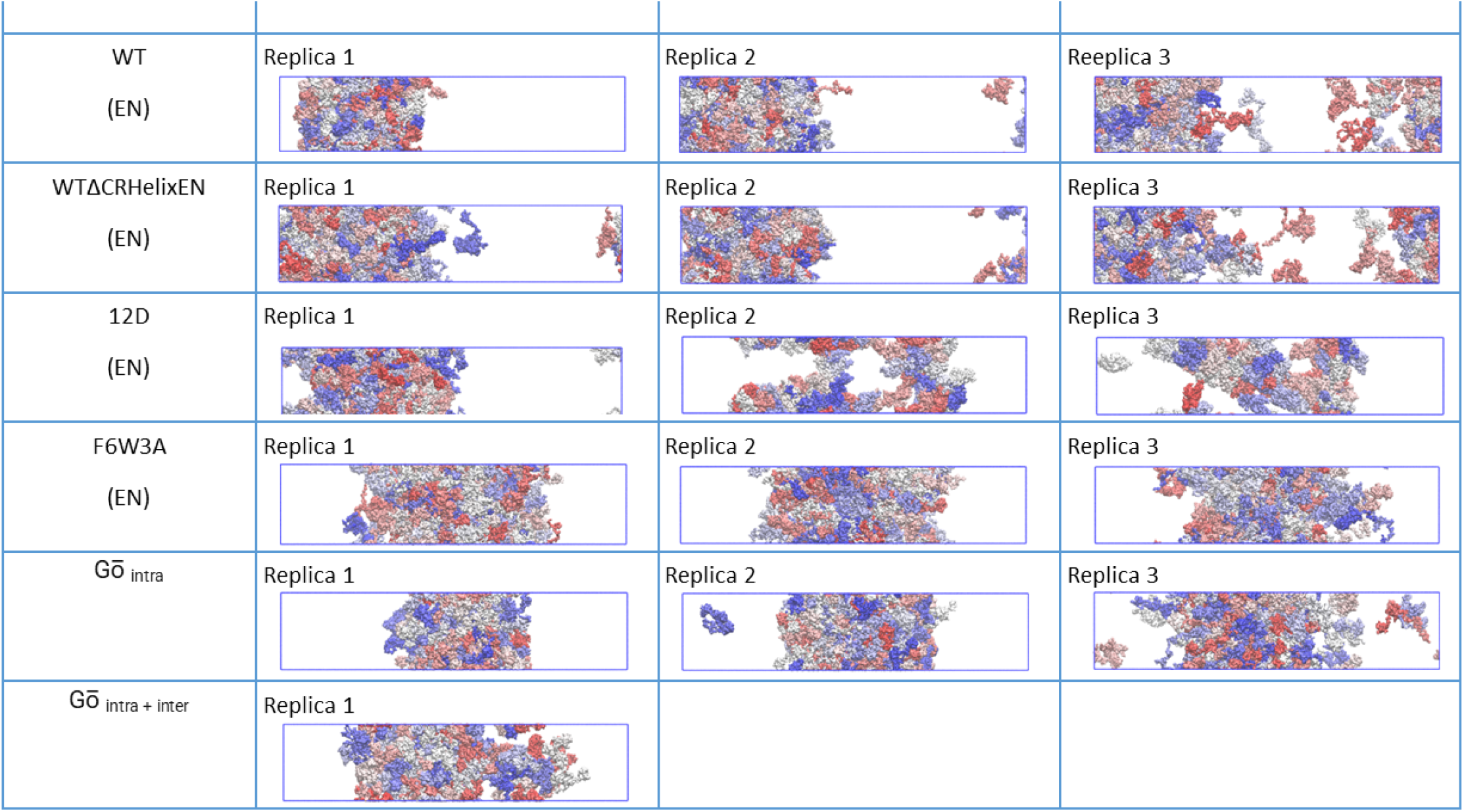
The initial configurations of the production simulation runs in slab geometry, water and ions are omitted for clarity.

**FIG. S5.**
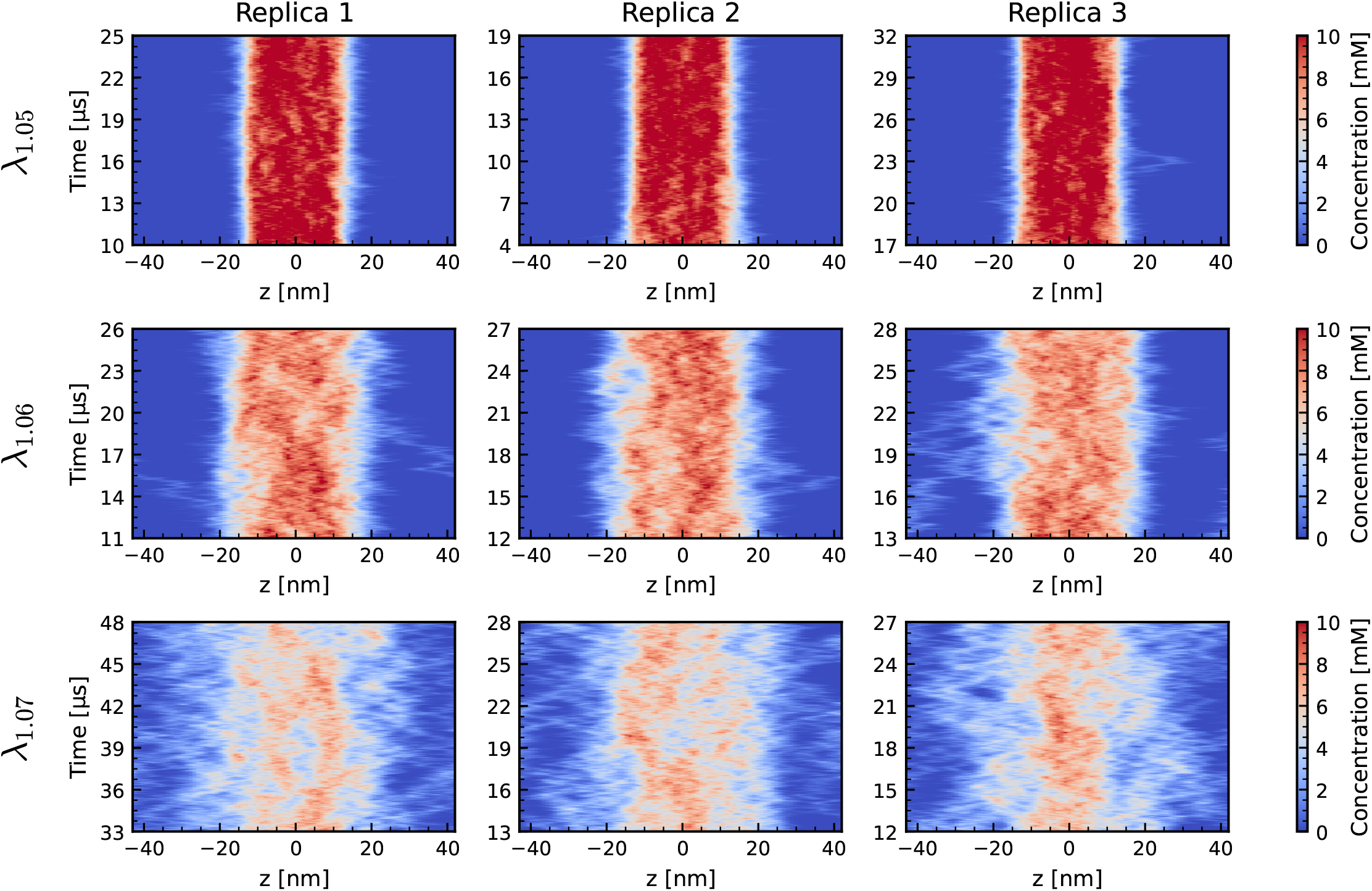
Time-dependent concentration histograms for full-length TDP-43 in Martini 3 Elastic Network (EN) simulations within slab geometry, shown for replicates at protein–water interaction strengths of *λ* = 1.05, *λ* = 1.06, and *λ* = 1.07.

**FIG. S6.**
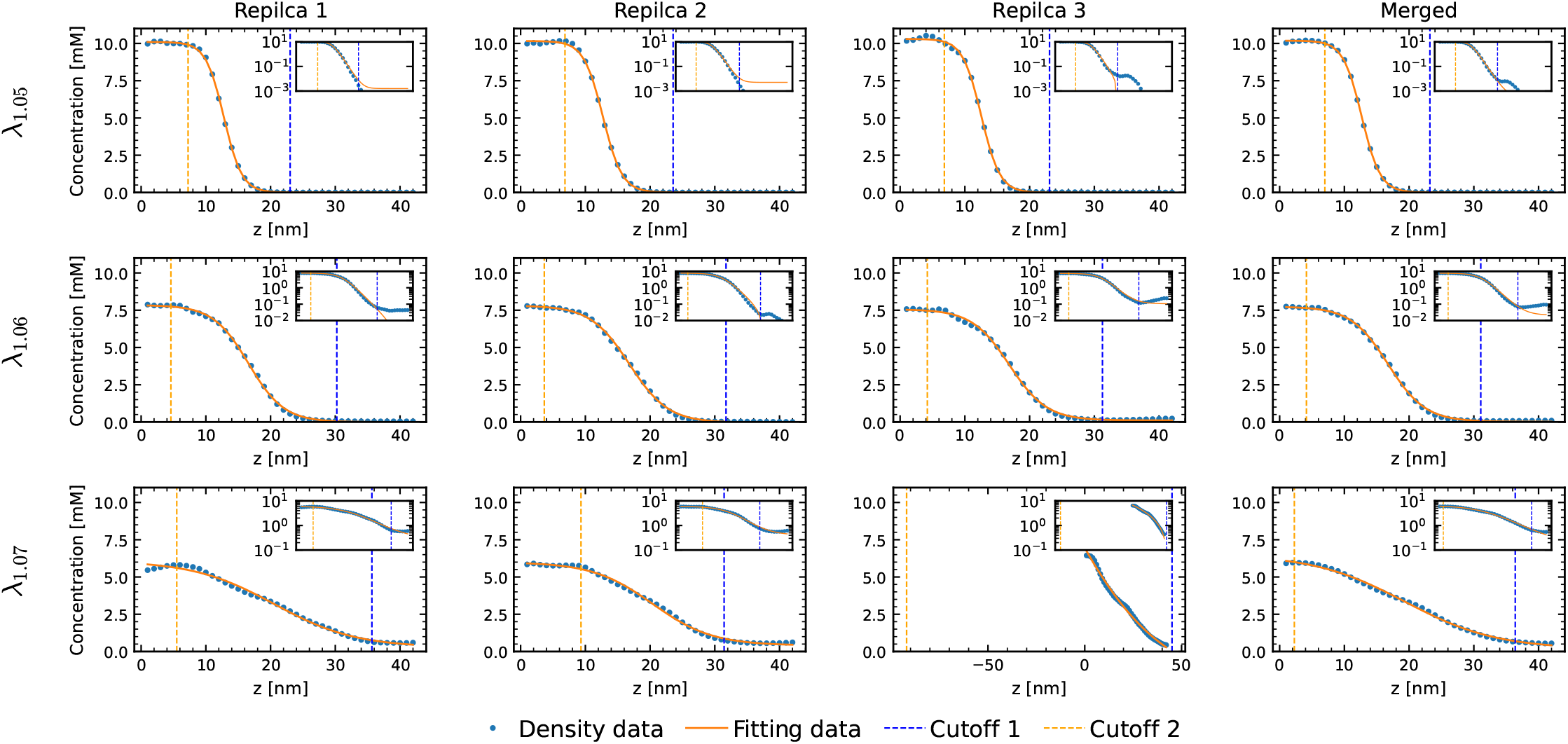
Averaged density histograms fitted to a hyperbolic tangent (tanh) function Eq. 2 to extract dense and dilute phase concentrations with the *Gō*_intra_ model at *λ* = 1.05, *λ* = 1.06 and *λ* = 1.07 in EN model.

**FIG. S7.**
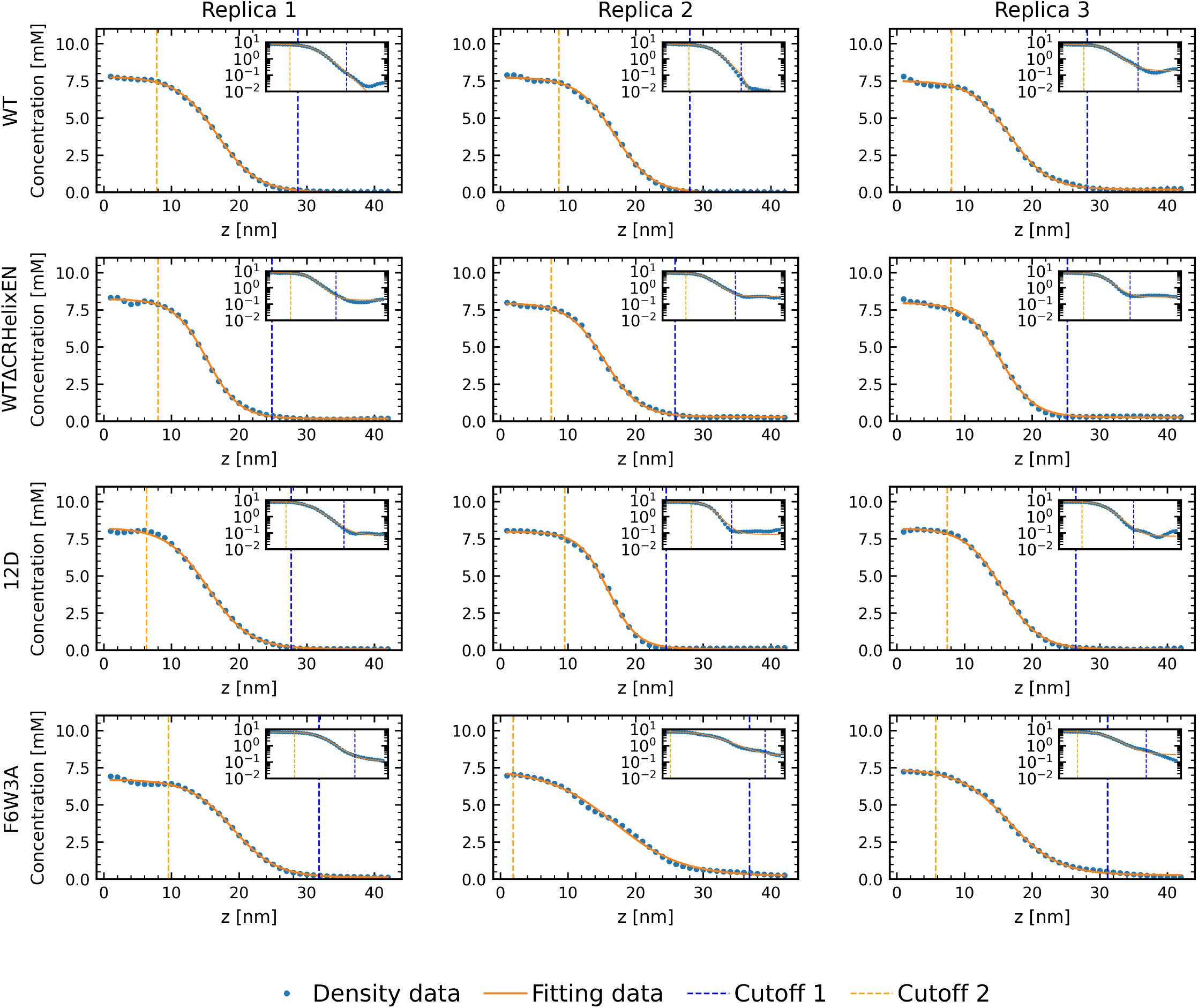
Averaged density histograms fitted to a hyperbolic tangent (tanh) function Eq. 2 to extract dense and dilute phase concentrations under the EN model at *λ* = 1.06 of TDP-43 and its mutants.

**FIG. S8.**
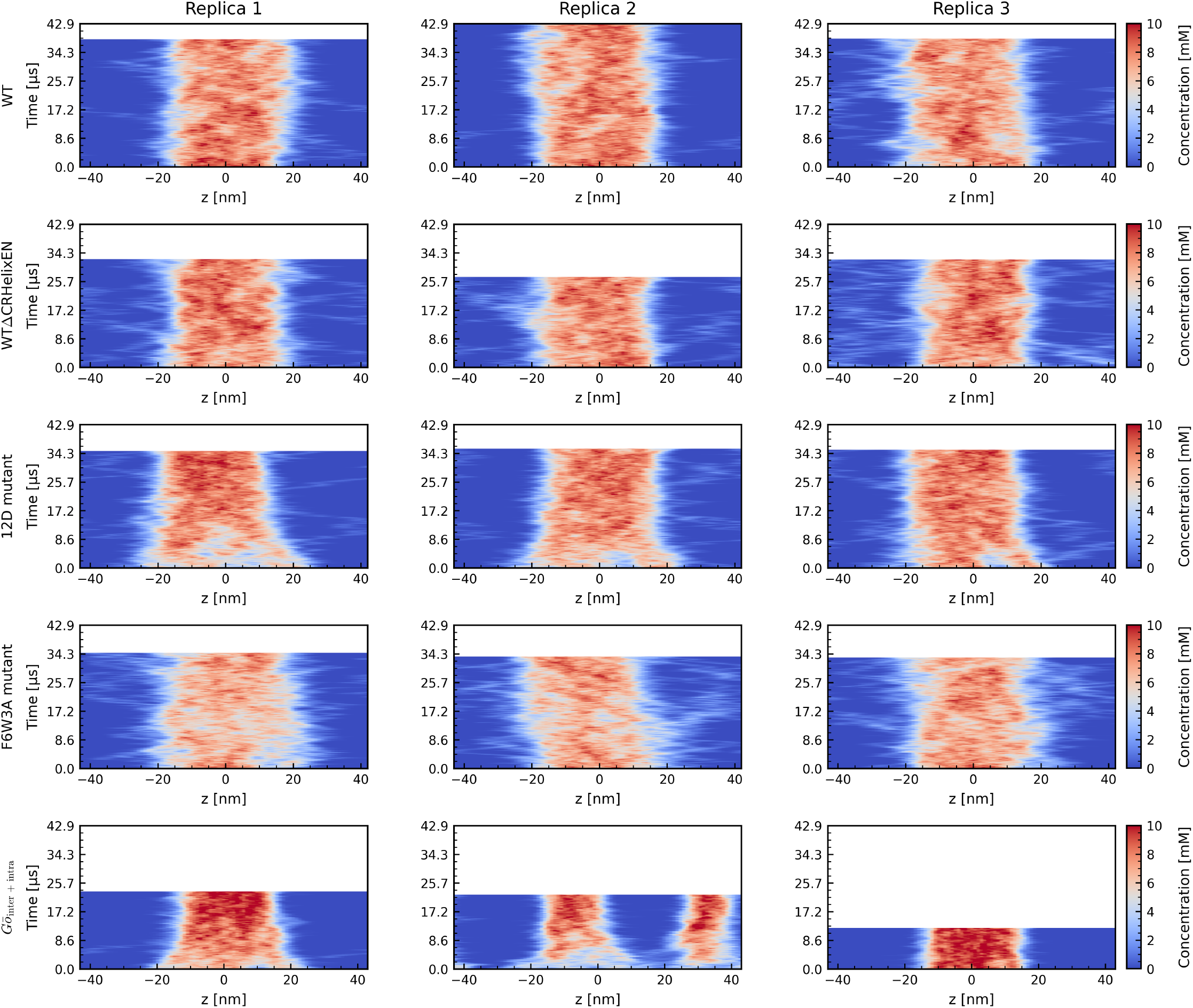
Time-dependent density histograms along the Z-axis across replicates of condensates for the EN model at *λ* = 1.06 of TDP-43 and its mutants and *Gō*_inter + intra_ model with *λ* = 1.06.

**FIG. S9.**
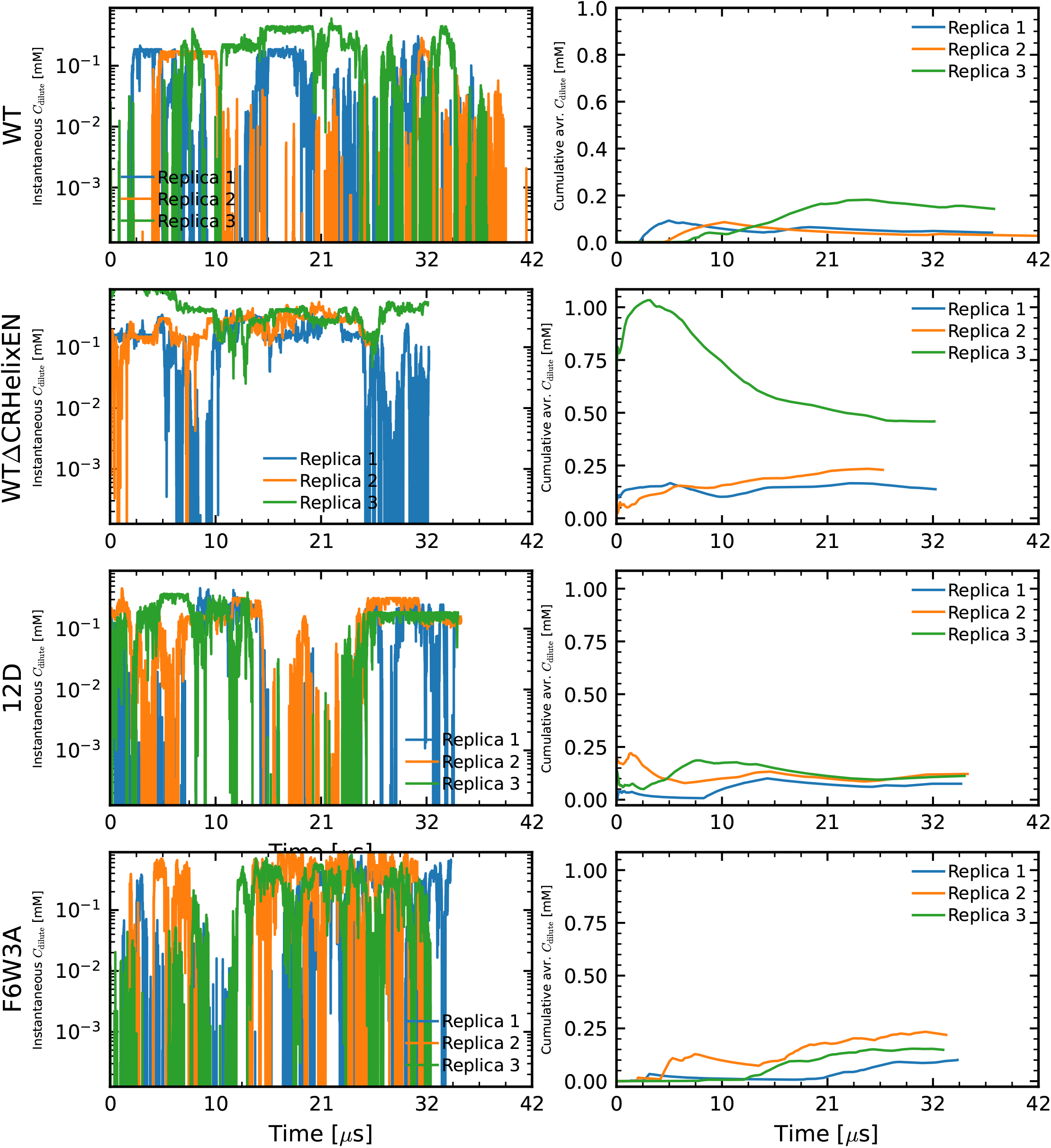
Analysis of dilute phase concentration (*c*_dilute_) using the *λ* = 1.06 EN model. (Left) Instantaneous *c*_dilute_ measurements for three independent replicas. (Right) Cumulative average of *C*_dilute_ over time.

**FIG. S10.**
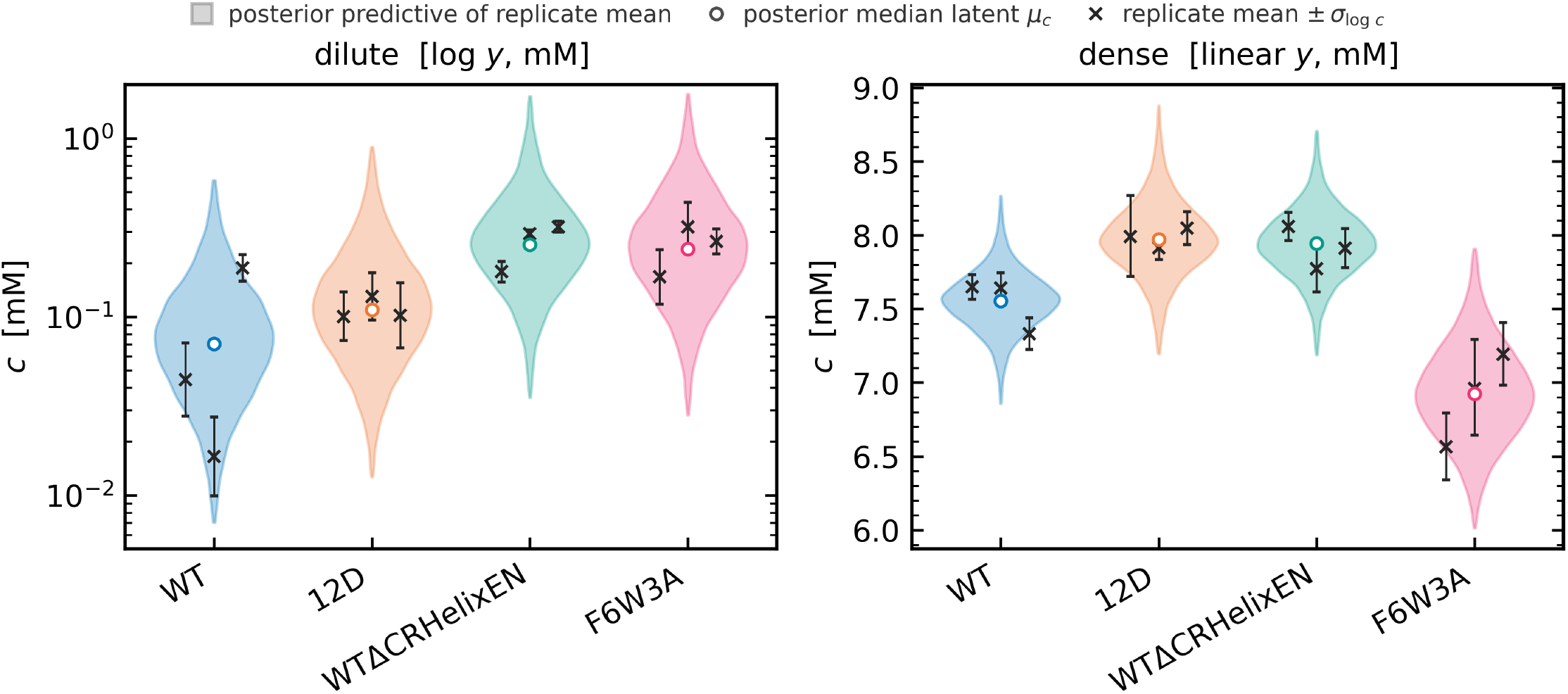
Posterior predictive checks of phase concentrations for TDP-43 variants. Posterior predictive distributions of replicate-mean concentrations from the Bayesian model for the dilute (left, logarithmic y-axis) and dense (right, linear y-axis) phases. Violin plots show posterior predictive distributions, circles indicate posterior median latent concentrations, and crosses represent replicate observations with asymmetric uncertainties derived from *σ*_log *c*_. Concentrations are reported in mM.

**FIG. S11.**
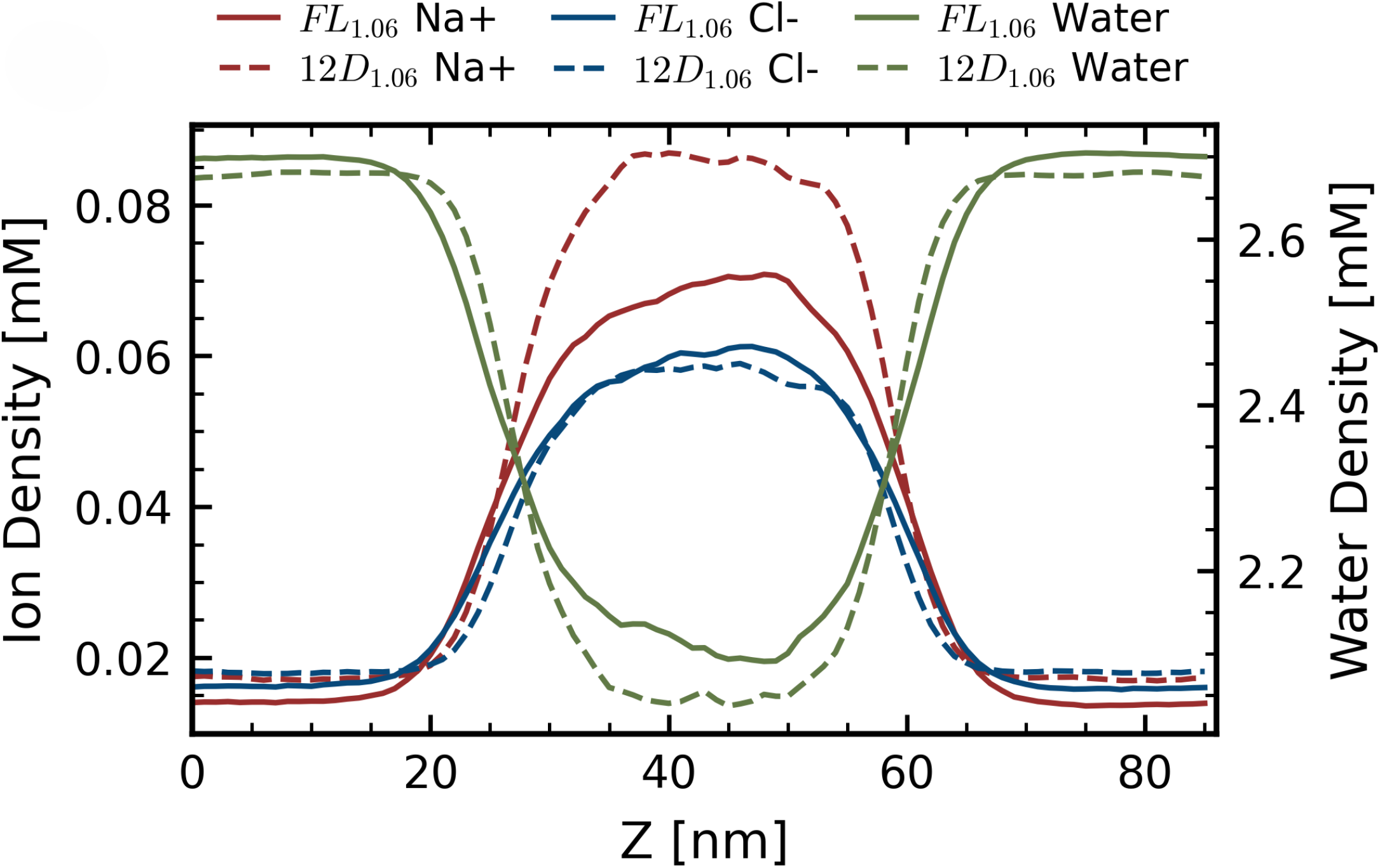
Ion and water density profiles along the z-axis for WT TDP-43 and the 12D mutant, highlighting differences in ion recruitment and local hydration within the condensate.

**FIG. S12.**
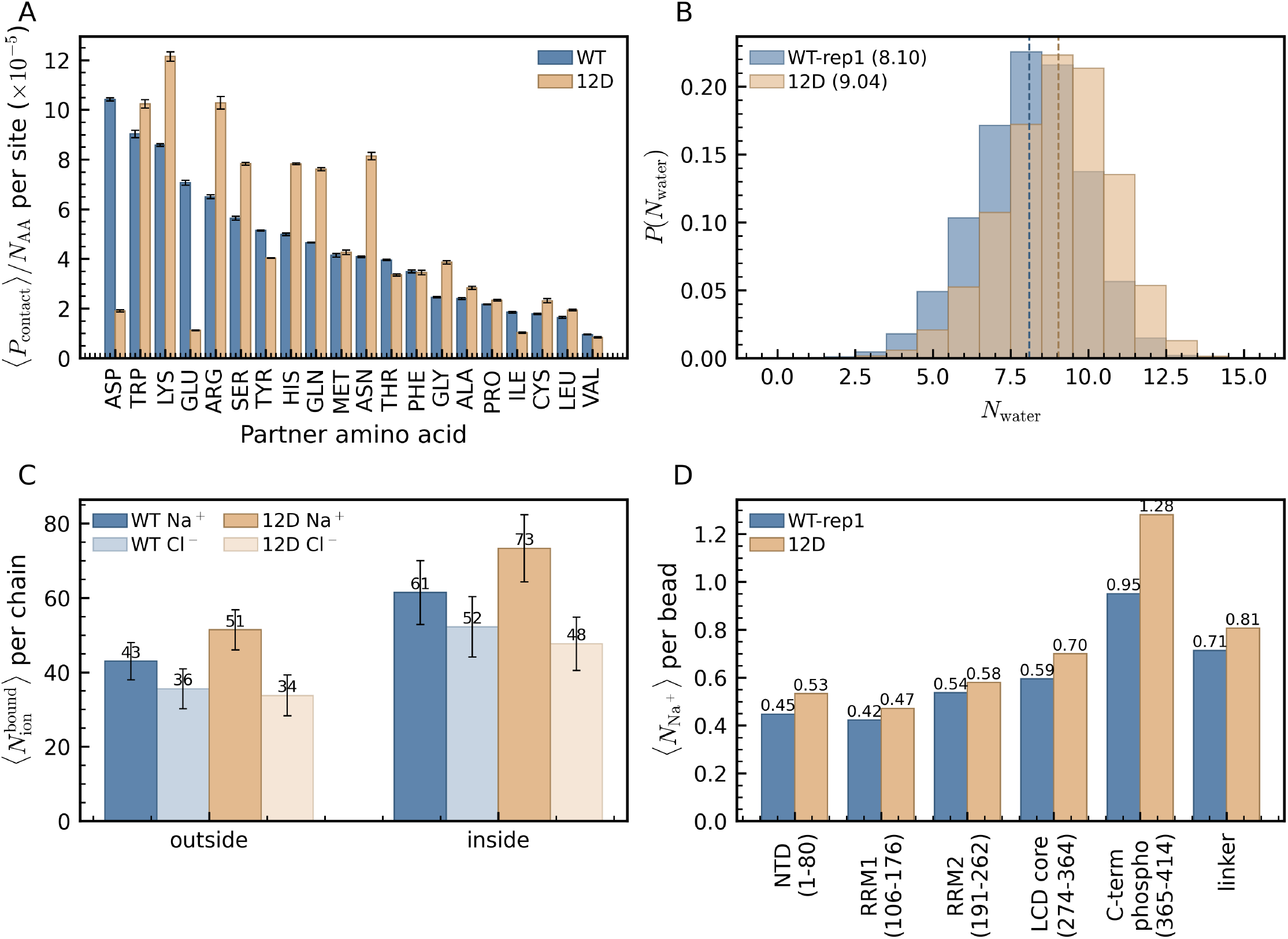
Effects of 12 phosphomimetic Ser→Asp mutations on contacts, water and ion binding. (A) ⟨*N*_partner_⟩ is the mean number of partner residues of type AA that are within 6Å of a Ser→Asp mutation site. All beads of the two amino acids are considered when evaluating whether a contact is formed. *N*_AA_ is the abundance of a given residue in the WT and 12D sequence. (B) The number of Martini water beads within a cut-off of 6.5 Å. of any bead of a phospho-site, sampled per chain × site × frame in the dense phase (*n* = 192,060 for WT-rep1, *n* = 106,392 for 12D; dashed lines mark the means. 8.10 vs. 9.04 W beads; 1 W bead ≡ 4 H_2_O). (C) Mean number of unique Na^+^ or Cl^−^ ions bound within 8 Å of any bead of a protein chain. Ion binding in dense and dilute phases is computed separately (dense, ≥200 inter-chain contacts within 10 Å) and outside (dilute, ≤100 contacts). (D) Cumulative first-shell Na^+^ coordination (*n*(*<* 8 Å)) per protein bead, averaged per TDP-43 region.

**FIG. S13.**
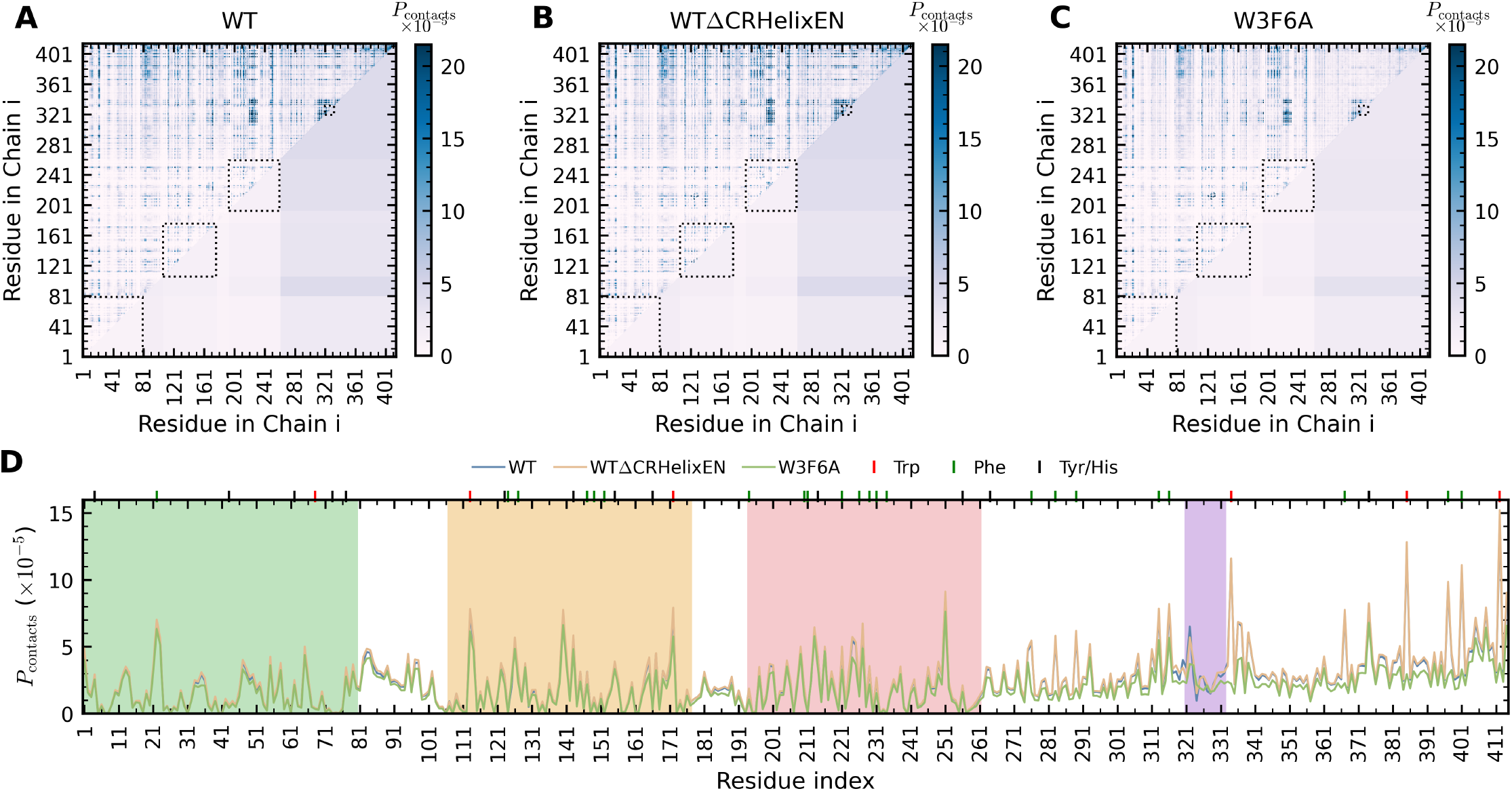
Intermolecular contacts within aromatic mutant condensates under EN simulations at *λ* = 1.06. (A–C). Inter-chain contact heatmaps for the WT protein, WTΔCRHelixEN and aromatic mutants F6W3A, respectively. Upper triangles are the residue-wise contact maps, lower triangles are domain-wise contact maps, residue- and domain-wise contact maps are sharing same colorbar. (D). Averaged intermolecular contact probability for each residue. Aromatic residues are indicated by colored vertical lines.

**FIG. S14.**
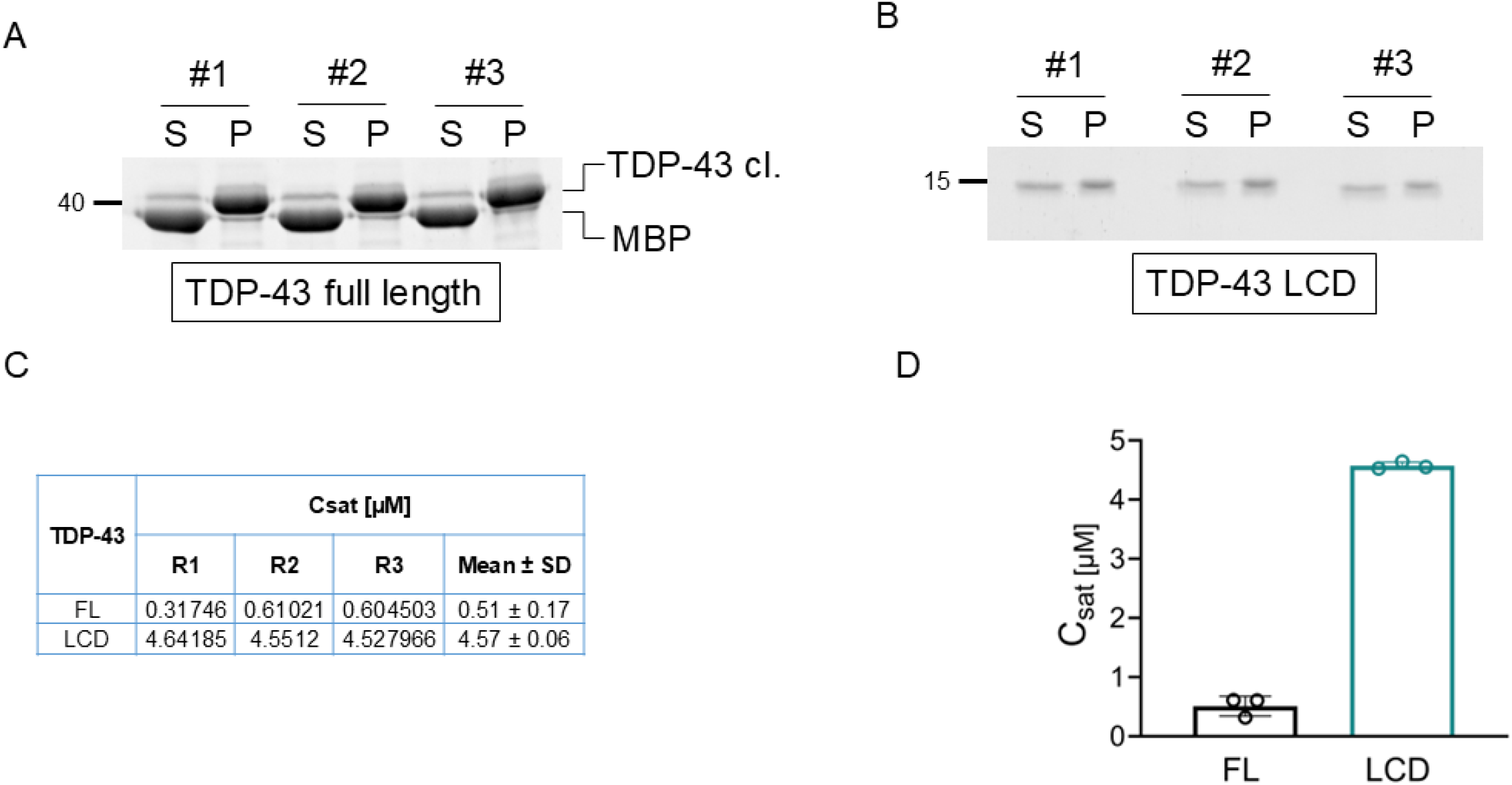
SyproRuby staining for sedimentation assay (at 10 µM protein concentration) to determine Csat of TDP-43 full length (A) and LCD (B). One representative replicate with three technical replicates each is shown. (C) *C*_*sat*_ values obtained from individual independent replicates together with the mean ± SD. (D) Calculated *C*_*sat*_ as mean of three indpendent replicates ± SD is shown for TDP-43 full length and LCD, respectively.

**FIG. S15.**
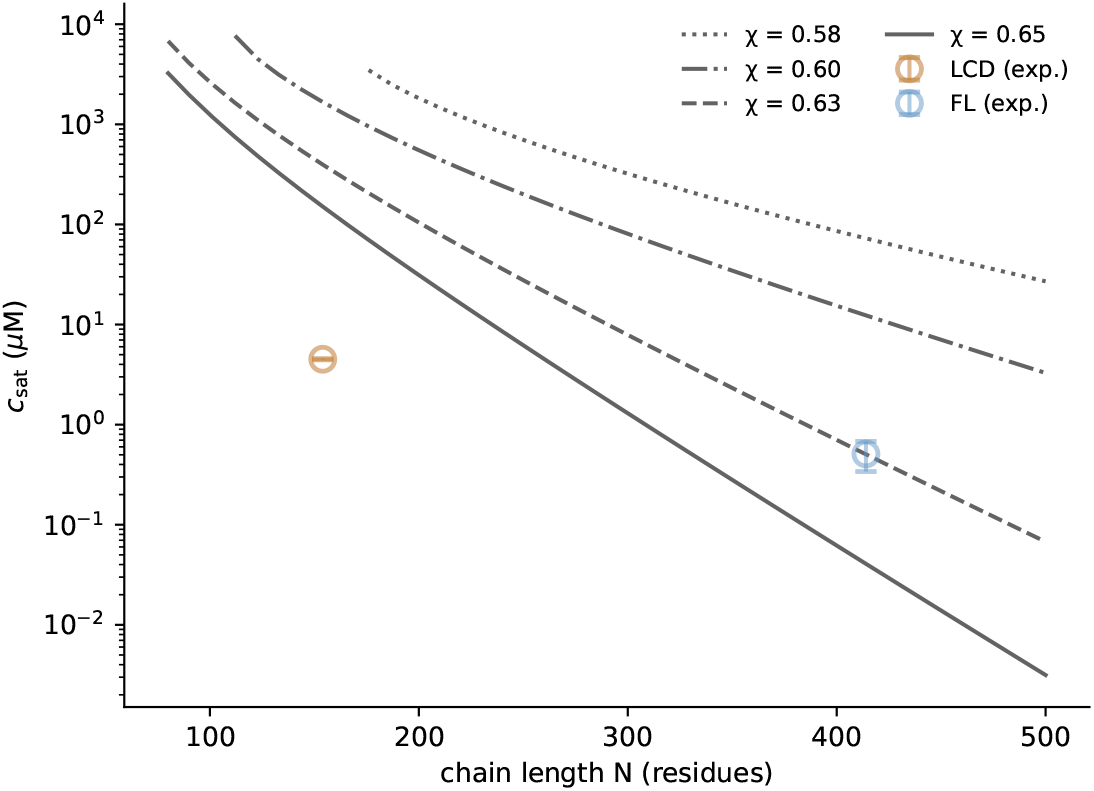
Comparison of *c*_dilute_ of full-length and LCD TDP-43 and chain length.

**FIG. S16.**
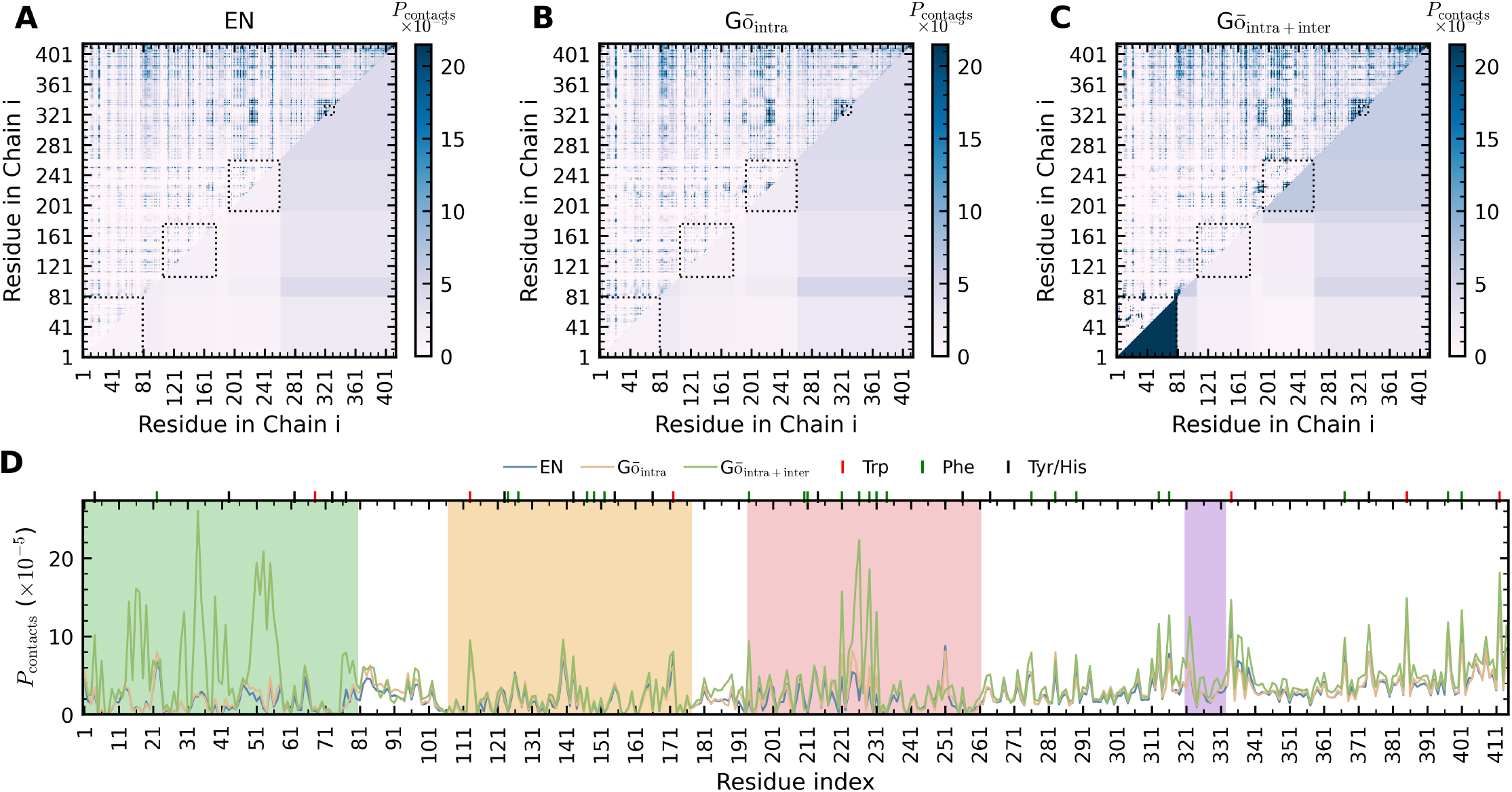
Inter-chain contacts within TDP-43 condensates under the EN, *Gō*_intra_, and *Gō*_intra+inter_ models at *λ* = 1.06. (A–C). Inter-chain contact heatmaps for the EN, *Gō*_intra_, and *Gō*_intra+inter_ models, respectively. Upper triangles show residue-wise contact maps, while lower triangles display domain-wise contact maps, calculated by summing the contact probabilities and averaging over the total number of residues within each domain. Residue- and domain-wise contact maps share the same colorbar. (D). Averaged inter-molecular contact probability of each residue across the three models: EN in blue, *Gō*_intra_ in orange, and *Gō*_intra+inter_ in dark green. Aromatic residues are indicated by colored vertical lines.

**FIG. S17.**
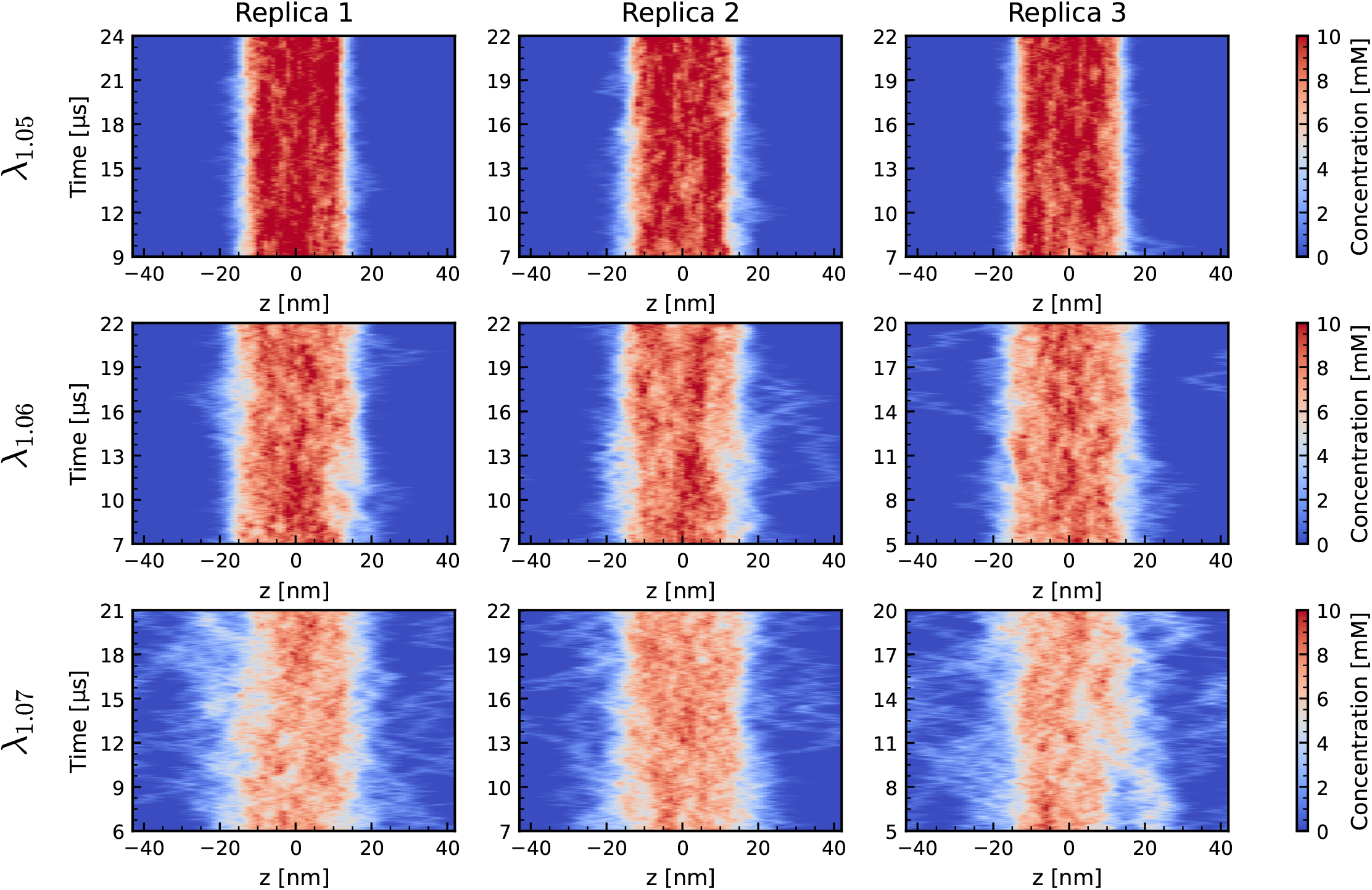
Time-dependent density histograms along the Z-axis across replicates of condensates for the *Gō*_intra_ model at *λ* = 1.05, *λ* = 1.06, and *λ* = 1.07 in *Gō*_intra_ model.

**FIG. S18.**
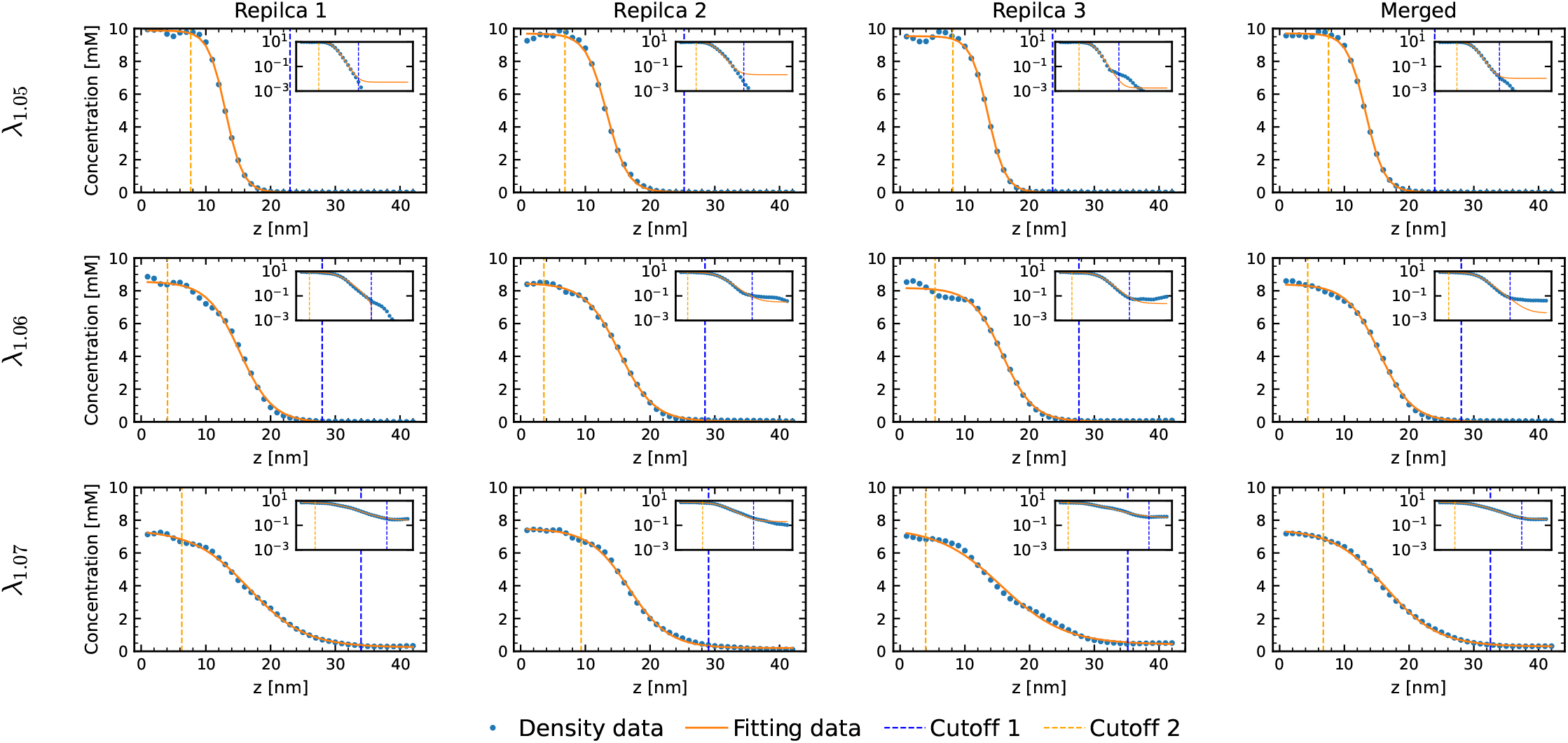
Averaged density histograms fitted to a hyperbolic tangent (tanh) function Eq. 2 to extract dense and dilute phase concentrations under the *Gō*_intra_ model at *λ* = 1.05, *λ* = 1.06 and *λ* = 1.07 in *Gō*_intra_ model.

**FIG. S19.**
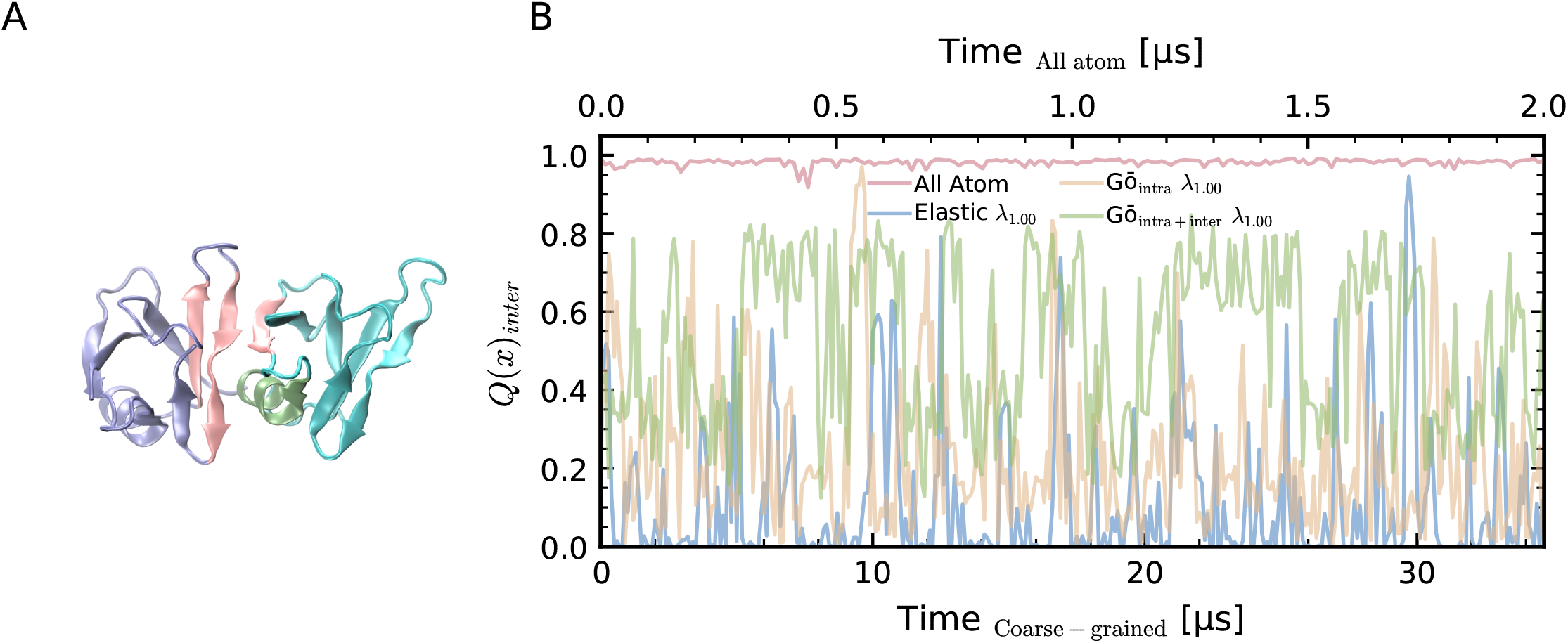
NTD dimerization. (A). Crystal Structure of the NTD-Dimer (PDB: 5MDI). The dimer is stabilized by complementary electrostatic interactions between the negatively charged head of one chain (right; *β*-sheet in pink and *α*-helix in green) and the positively charged tail of the partner chain (left; *β*-sheet in pink). (B). Fraction of native inter-chain contacts observed in all-atom and coarse-grained simulations of the NTD homodimer.

**FIG. S20.**
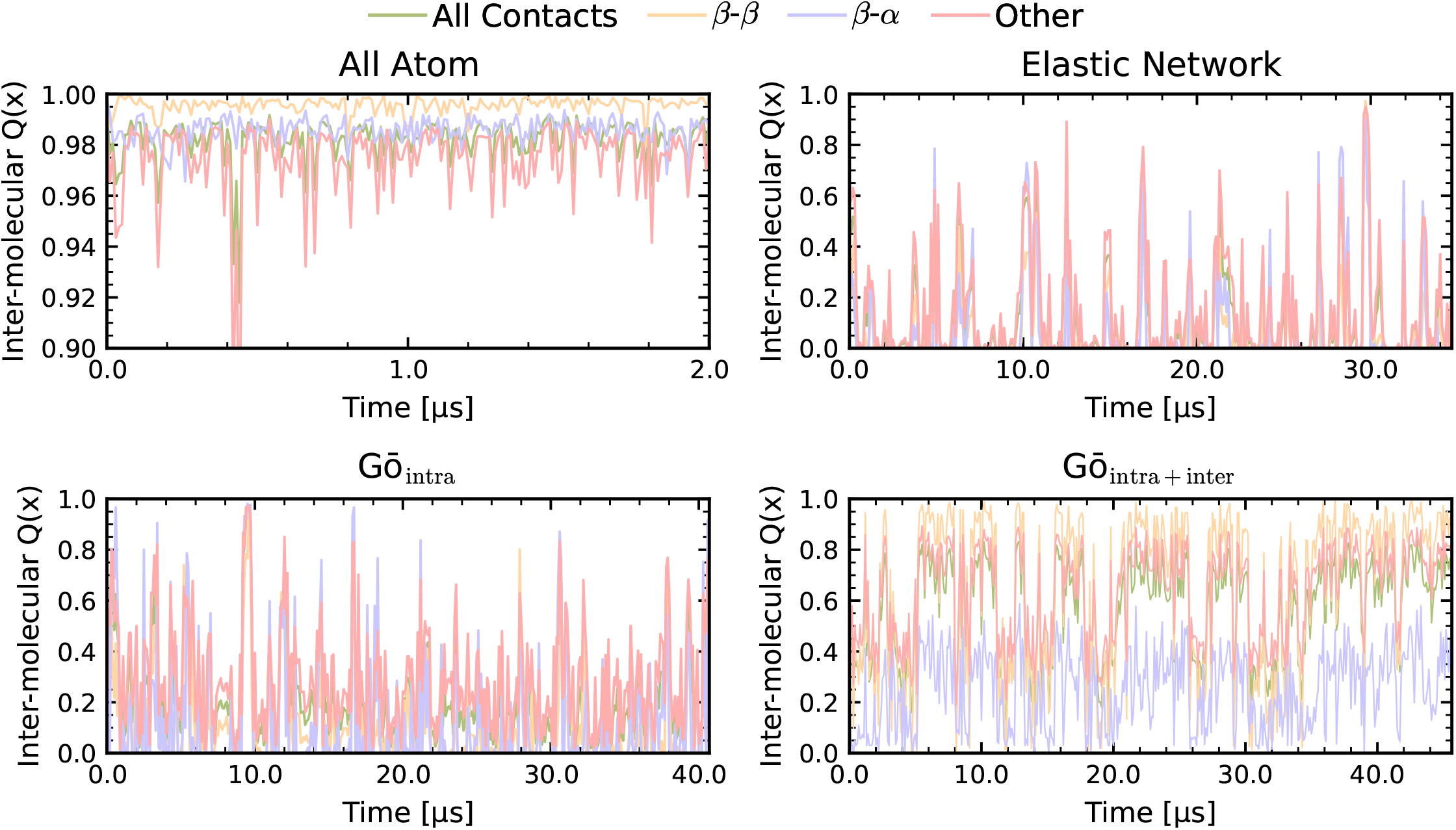
Decomposition of native inter-chain contacts in NTD homodimer. Breakdown of *Q*_*x*_ into individual interface components: *β*-*β, β*-*α*, and other contacts for (A) AA, (B) EN, (C) *Gō*_intra_, and (D) *Gō*_intra+inter_ models.

**FIG. S21.**
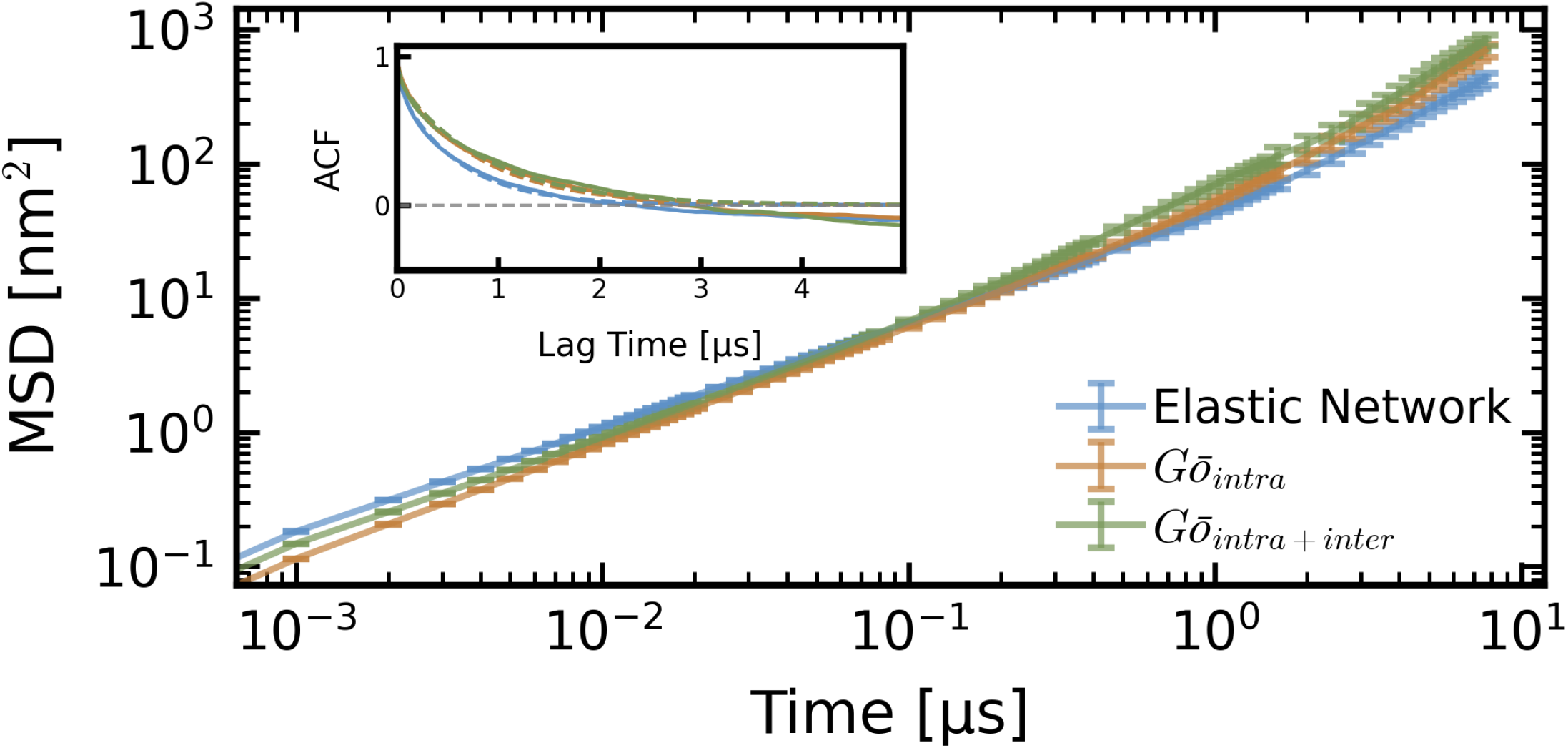
Mean square displacement (MSD) of TDP-43 in three models with replicas, the inset shows end-to-end distance autocorrelations for each model at *λ* = 1.06.

**FIG. S22.**
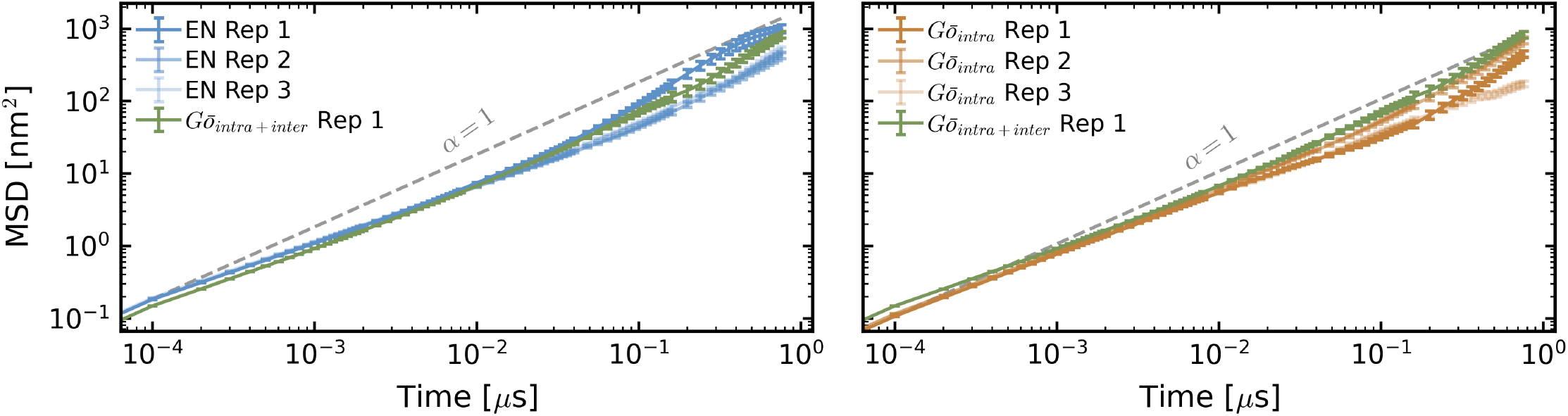
Comparison of MSD for full-length TDP-43. (Left) Comparison between the EN model and the *Gō*_intra+inter_ model. (Right) Comparison between the *Gō*_intra_ and *Gō*_intra+inter_ models, both simulated at a scaling factor of *λ* = 1.06. Dashed lines indicate a slope of *α* = 1 as a reference for normal diffusion.

**FIG. S23.**
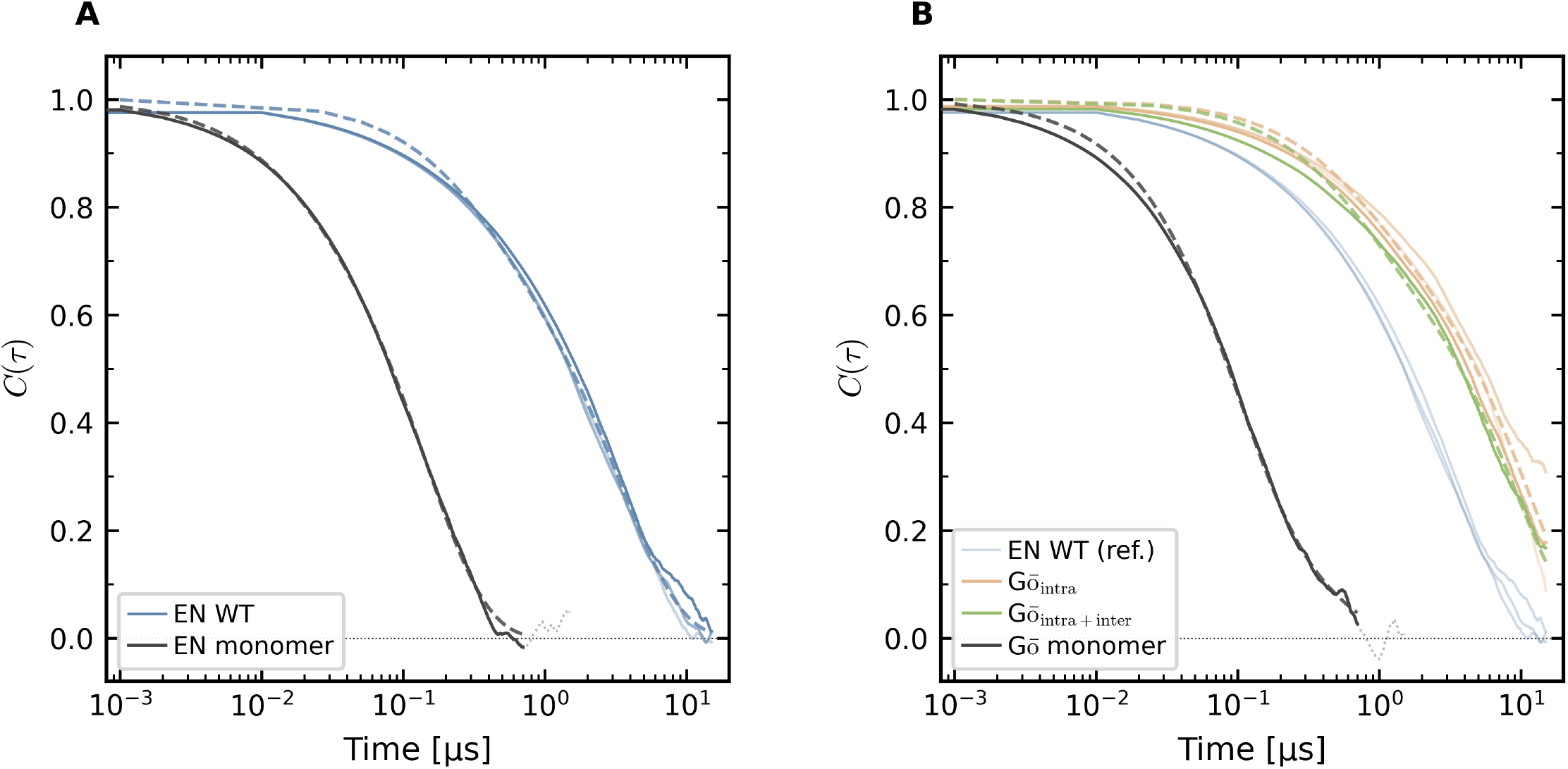
End to end distance autocorrelation function decays and bi-exponential fit analysis of dilute and dense phases. Solid lines display the normalized autocorrelation functions *C*(*τ*), and dashed lines represent corresponding bi-exponential fits, *C*(*τ*) = *A* · exp(−*τ/τ*_1_) + (1 − *A*) · exp(−*τ/τ*_2_). (A) Comparison of relaxation dynamics between the EN model in the dense phase (blue line; fast relaxation *τ*_1_ = 341 ns, slow relaxation *τ*_2_ = 3.45 µs, and mean relaxation ⟨*τ* ⟩ = 2.76 µs) and an isolated EN single chain in the dilute phase (black line; fast relaxation *τ*_1_ = 16 ns, slow relaxation *τ*_2_ = 146 ns, and mean relaxation ⟨*τ* ⟩ = 0.131 µs). (B) Comparison of relaxation dynamics for the single chain in the dilute phase using the Gō model (black line; black line; fast relaxation *τ*_1_ = 92 ns, slow relaxation *τ*_2_ = 402 ns, and mean relaxation ⟨*τ* ⟩ = 0.174 µs) evaluated against two reference Gō-type models: the Gō _intra_ model (fast relaxation *τ*_1_ = 715 ns, slow relaxation *τ*_2_ = 10.6 µs, and mean relaxation ⟨*τ* ⟩ = 8.50 µs) and the Gō_intra + inter_ model (green line; fast relaxation *τ*_1_ = 645 ns, slow relaxation *τ*_2_ = 8.95 µs, and mean relaxation ⟨*τ* ⟩ = 6.94 µs). The EN WT dense-phase curve is provided for reference.

**FIG. S24.**
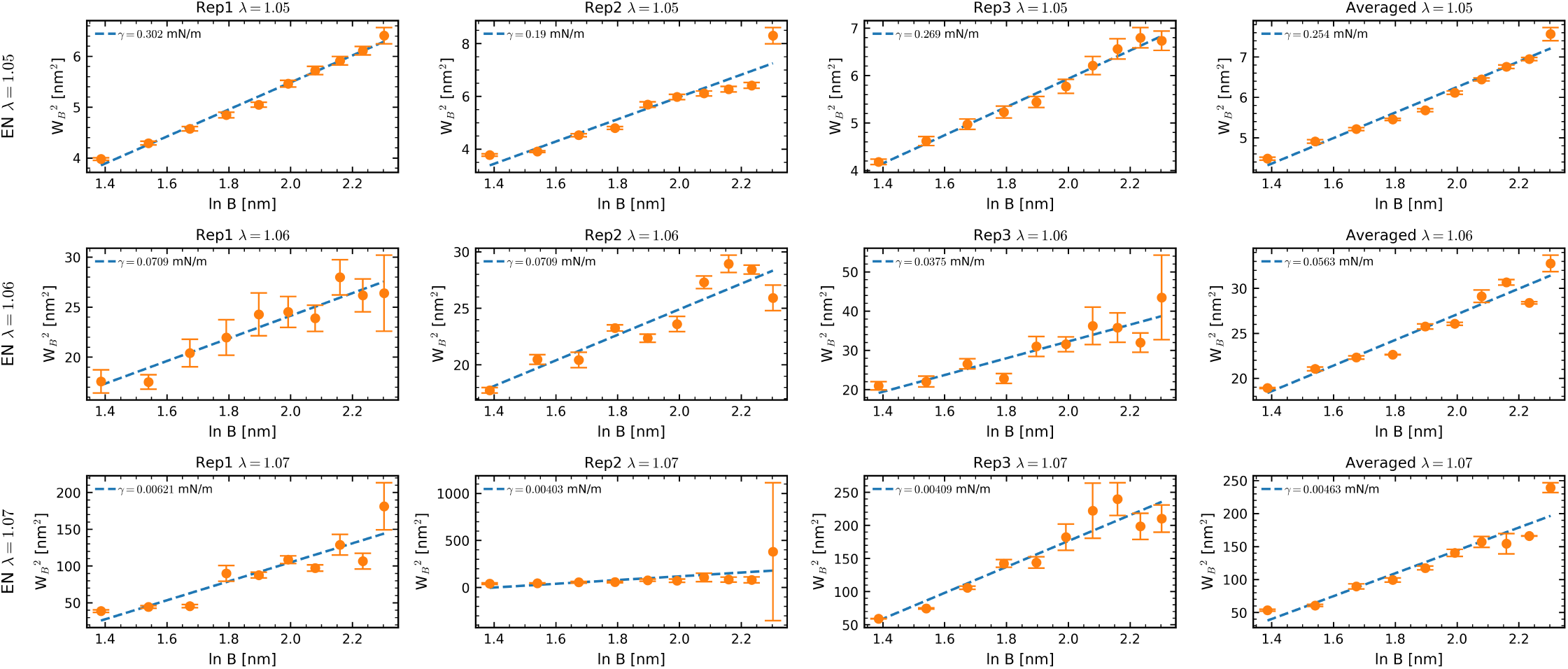
Interfacial width *W*^2^ as a function of block size *B* in EN model. The interface region is divided into rectangular segments, and the interfacial tension *γ* is determined using 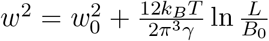, where *W*_0_ is the intrinsic interfacial width, *k*_*B*_ is the Boltzmann constant, *T* is the temperature, and *L* is the simulation box size.

**FIG. S25.**
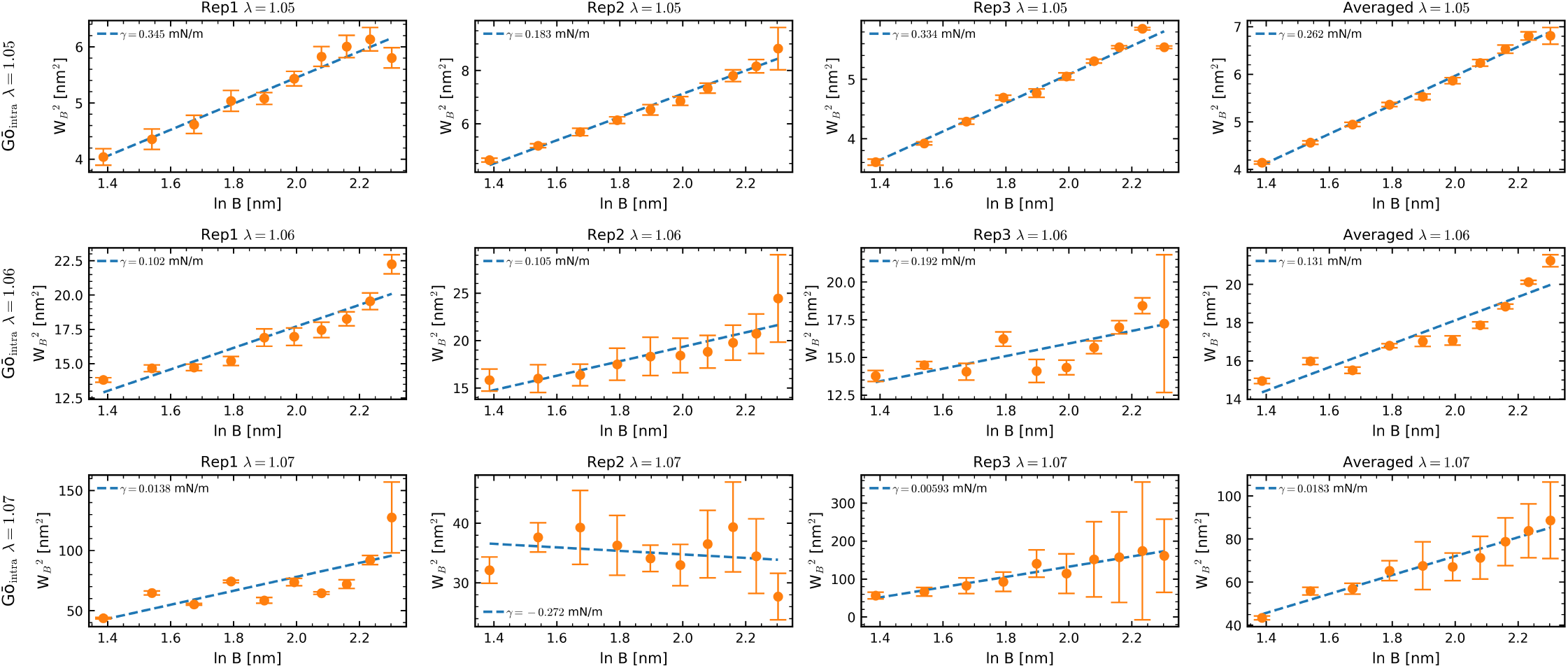
Interfacial width *W*^2^ as a function of block size *B* in *Gō*_intra_ model.

**FIG. S26.**
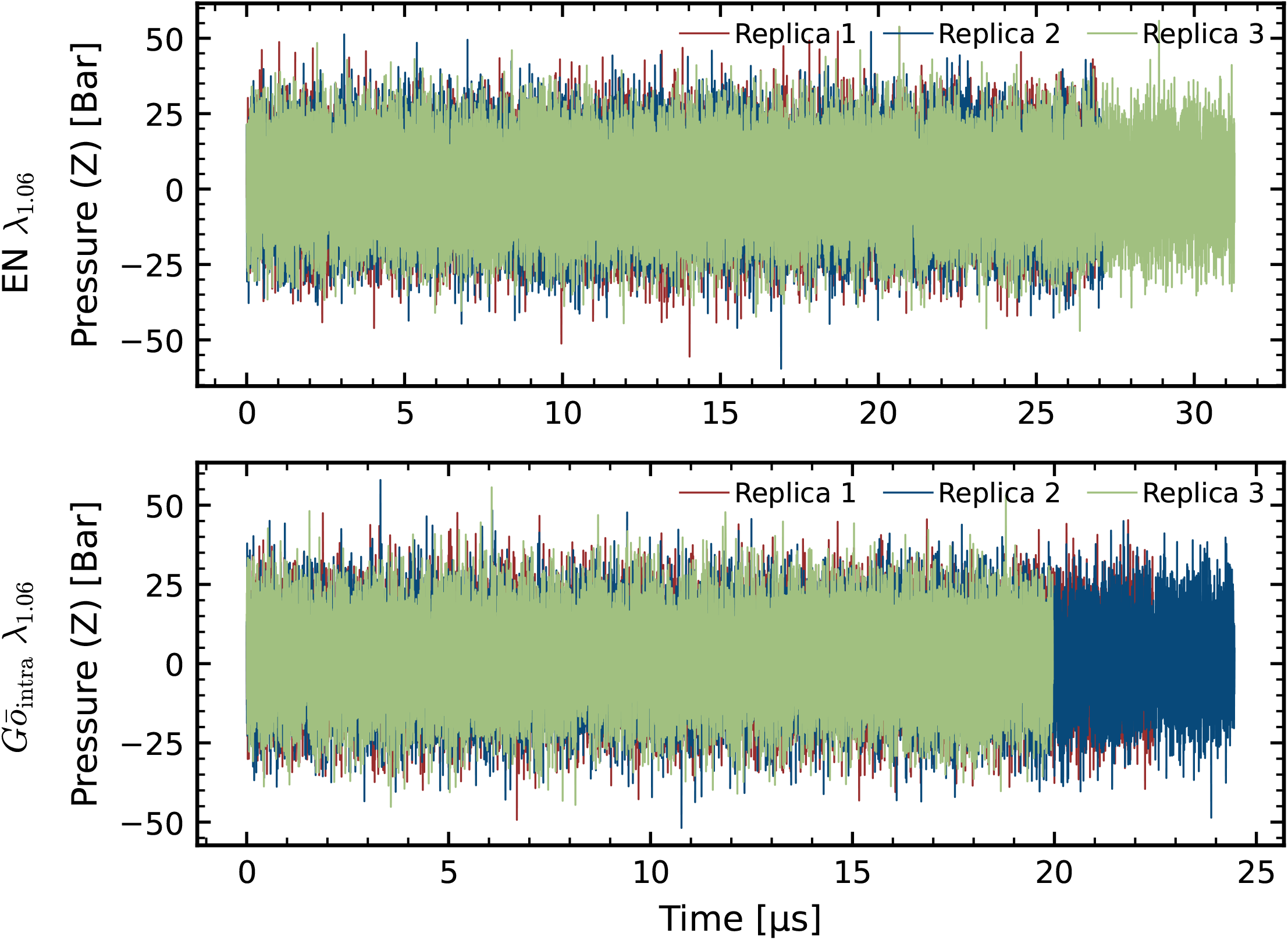
Evolution of the box pressure along the *z*-axis over time for three replicas of Martini 3 Elastic Network (EN) (top) and *Gō*_intra_ (bottom) simulation systems in slab geometry, with protein–water interaction strength *λ* = 1.06.

**FIG. S27.**
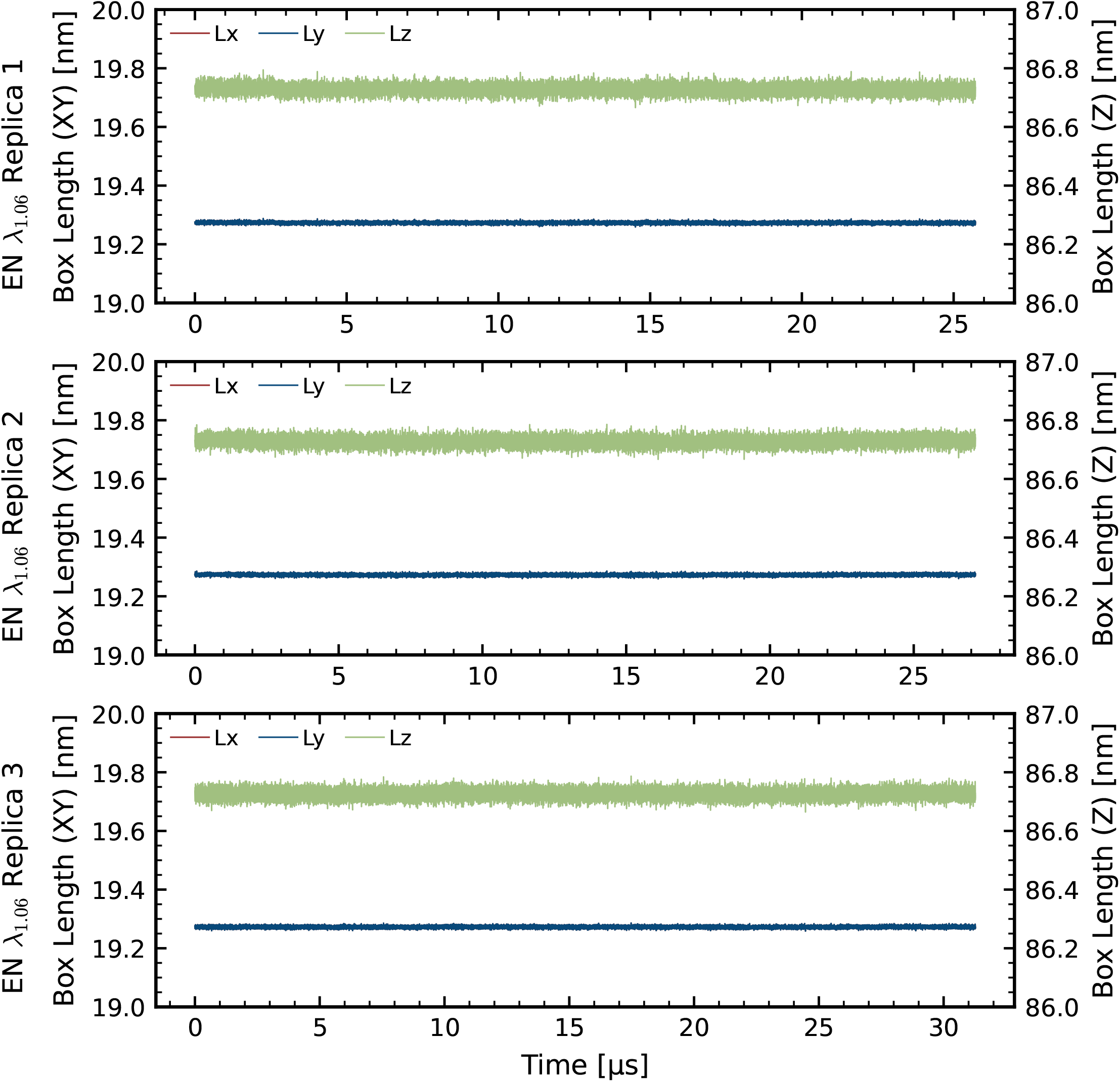
Box length evolution over time for three replicas of the Martini 3 Elastic Network (EN) simulations system in the slab geometry with protein–water interaction strengths of *λ* = 1.06. The XY dimensions (Lx, Ly) are shown on the left y-axis, while the Z dimension (Lz) is shown on the right y-axis.

**FIG. S28.**
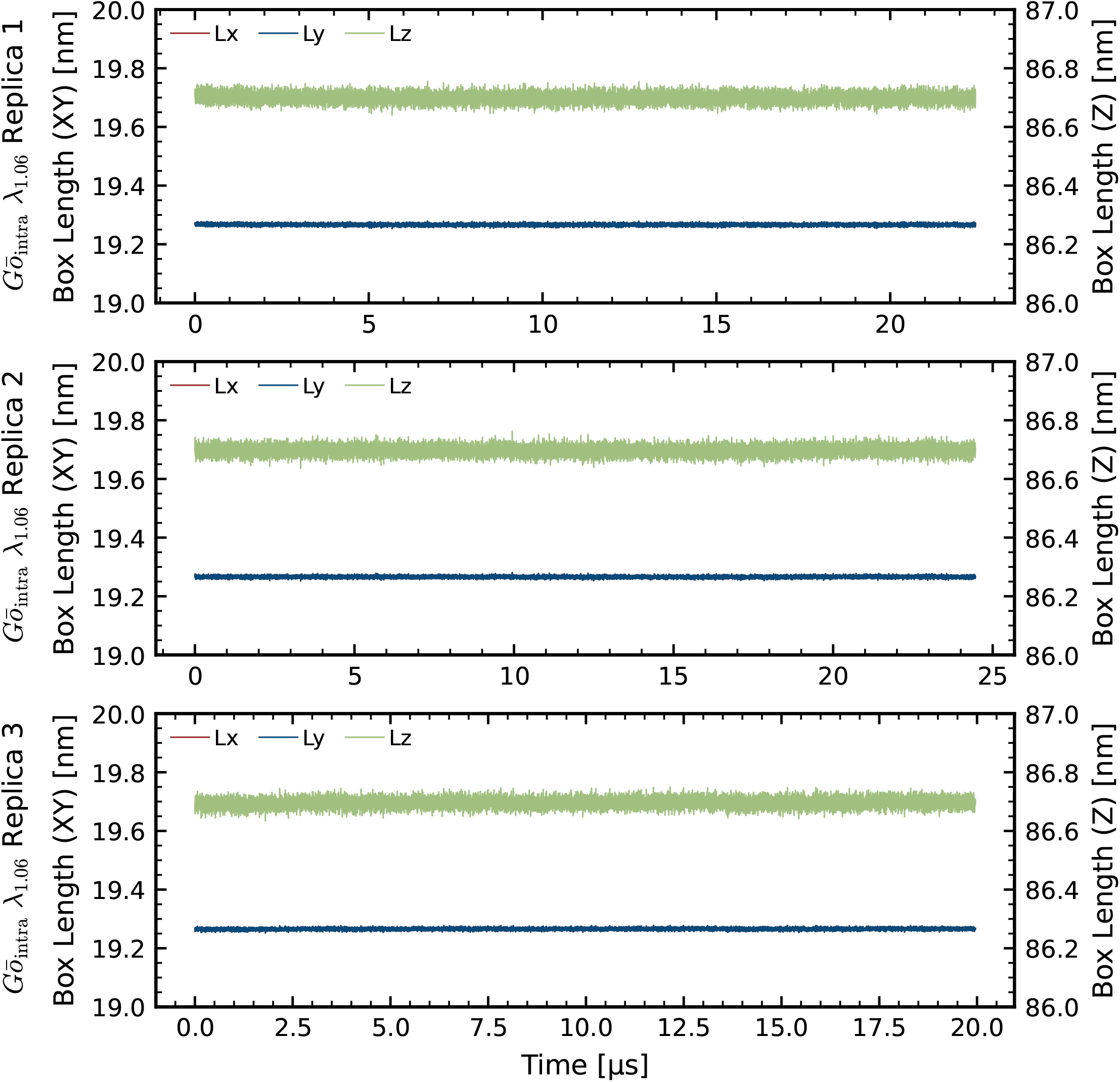
Box length evolution over time for three replicas of the Martini 3 *Gō*_intra_ simulations system in the slab geometry h protein–water interaction strengths of *λ* = 1.06. The XY dimensions (Lx, Ly) are shown on the left y-axis, while the Z dimension (Lz) is shown on the right y-axis.

## Notes

### Competing Interest Statement

The authors have declared no competing interest.

### Summary of Updates

this version has updates figure2 in the main text

